# Phylogenomics and comparative genomics of the genus *Erwinia* reveal taxonomic inconsistencies and evolutionary diversification

**DOI:** 10.64898/2026.08.06.743344

**Authors:** Nimisha Maurya, Shefali Dobhal, George Sundin, Brendan Rodoni, James P. Stack, Mohammad Arif

## Abstract

The genus *Erwinia* comprises a diverse group of bacteria associated with plants, insects, and the environment, including several economically important phytopathogens. The genus has been revised taxonomically many times, yet a thorough and genome-wide assessment of its evolutionary relationships and genomic diversity has been lacking. In this research, we carried out an extensive phylogenomic and comparative genomic analyses of the genus *Erwinia* using 104 genomes including historically important strains. Genome-wide analyses integrating average nucleotide identity (ANI), digital DNA–DNA hybridization (dDDH), core-genome phylogenomics, pan-genome analysis, and comparative genomics resolved evolutionary relationships across the genus and identified multiple taxonomic inconsistencies. The pan-genome analysis revealed a relatively small core genome alongside an extensive accessory genome, underscoring the substantial genomic plasticity and ongoing diversification within the genus. The comparative analyses further showed pronounced lineage-specific variation in secretion systems, exopolysaccharide biosynthetic loci, flagellar gene clusters, genomic islands, prophages, and iron acquisition systems, suggesting that virulence-associated determinants have evolved through differential gene gain, loss, and conservation across distinct lineages, thereby facilitating host and ecological niche adaptation. This lineage-specific variation indicates that pathogenicity in the genus is not driven by a single conserved set of virulence determinants but instead reflects distinct combinations of virulence-associated genes. These findings refine the genomic framework of the genus *Erwinia*, provide evidence for taxonomic revision of several lineages, and improve our understanding of the evolutionary relationships, genomic diversification, and lineage-specific adaptations associated with host interactions and ecological specialization.

**Impact Statement:** This study provides the first comprehensive genome-wide phylogenomic framework for the genus *Erwinia*, integrating taxonomy, pan-genome diversity, virulence-associated determinants, and mobile genetic elements across all 18 currently recognized species. Analyses resolve evolutionary relationships, uncover multiple taxonomic inconsistencies, identify previously unrecognized species-level lineages, including a putative novel *Erwinia* species PL328 isolated from *Cornus florida* (dogwood), and reveal lineage-specific genomic features. These findings establish a valuable genomic foundation for future studies of *Erwinia* evolution, taxonomy, and plant-microbe interactions.

**Data Summary:** Genomes sequenced in this study were submitted to the NCBI database under the accession numbers: JCBCPT000000000

## Introduction

*Erwinia,* within the *Erwiniaceae* family, is a genus of Gram-negative, rod-shaped, motile bacteria first described by Winslow et al. [1]. The genus includes several important plant-associated species responsible for economically significant plant diseases, such as fire blight, shoot blight, bacterial wilt, stem necrosis, and soft rot, collectively causing major yield losses in fruit trees, vegetables, and ornamental crops worldwide [2–5]. These pathogens produce characteristic symptoms, including wilting, necrosis, water-soaked lesions, vascular discoloration, and tissue maceration, depending on host and pathogen involved [2,5,6,7].

The genus *Erwinia* has undergone substantial taxonomic revision since its initial description in the early 20th century. Analyses of nearly complete 16S rRNA gene sequences revealed that several taxa previously classified within *Erwinia*, including species now assigned to *Pectobacterium*, *Dickeya*, *Pantoea* and *Brenneria,* represented distinct phylogenetic lineages and were subsequently reclassified into separate genera [8]. Phylogenomic studies further refined its classification, leading to the placement of *Erwinia* within the family *Erwiniaceae* of the order Enterobacterales. According to the List of Prokaryotic Names with Standing in Nomenclature (LPSN) database [9] at https://lpsn.dsmz.de, 18 species are currently recognized within the genus as of November 5, 2025, as shown in **Figure 1**.

**Figure 1.**
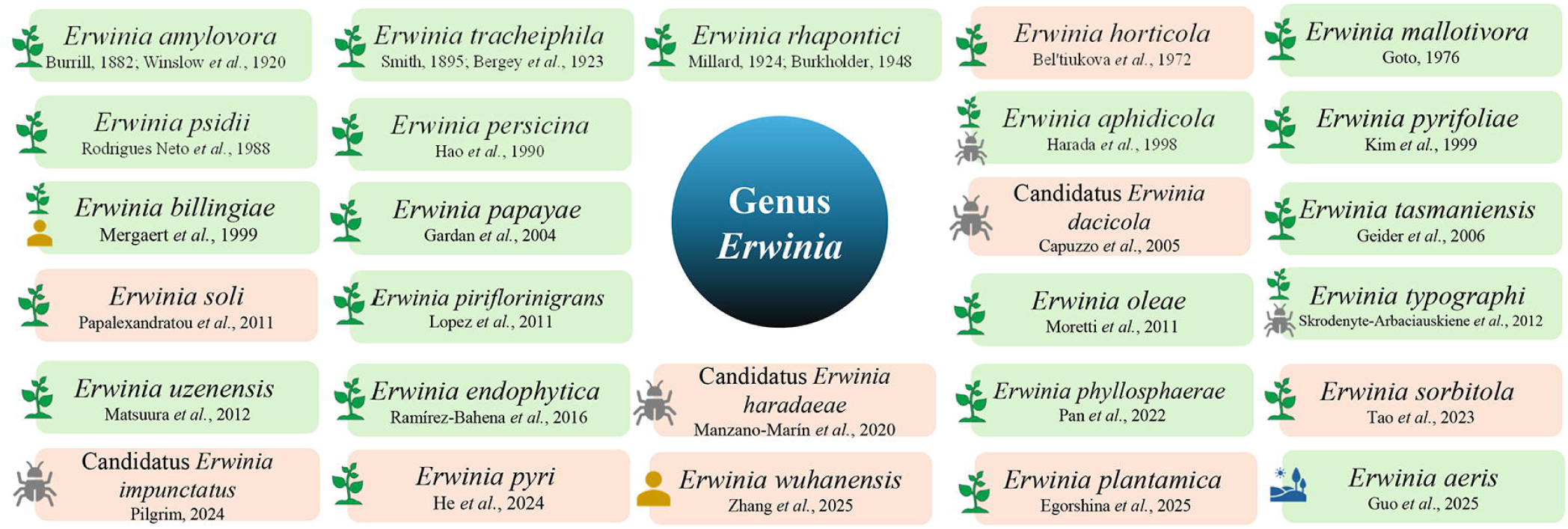
Overview of all validly published and not validly published species assigned to the genus *Erwinia*, based on the current taxonomic status in the List of Prokaryotic names with Standing in Nomenclature (https://lpsn.dsmz.de/). Species shown in green box represent validly published taxa, whereas those shown in orange box represent not validly published taxa. Icons indicate the major reported ecological association(s) of each species: plant (green plant icon), environmental (blue landscape icon), human (yellow human icon), and insect (gray insect icon).

The genus comprises both pathogenic and non-pathogenic plant-associated bacteria exhibiting broad ecological and host diversity. Among the pathogenic members, *E. amylovora* and *E. pyrifoliae* are well-known pathogens, causing fire blight and shoot blight in members of the Rosaceae [1,10]. Several other species, including *E. mallotivora*, *E. psidii*, *E. piriflorinigrans*, *E. uzenensis*, and *E. papaya,* infect a wide range of hosts such as *Mallotus japonicus*, guava, pear and papaya, producing diverse symptoms including leaf spots, blossom necrosis, shoot blight, and stem cankers [11–15]. Other pathogenic species, such as *E. persicina* and *E. rhapontici*, and the recently described *E. pyri*, have been associated with soft rot in vegetables and dieback in pear [5,16–19], whereas *E. tracheiphila* is the causal agent of bacterial wilt in cucurbits [20]. In contrast, several *Erwinia* species, such as *E. tasmaniensis*, *E. endophytica*, *E. oleae*, and *E. phyllosphaerae,* are considered as non-pathogenic and plant-associated endophytes [21–24]. Beyond plants, the genus also includes insect-associated species, such as *E. aphidicola*, *E. sorbitola* and *E. typographi,* highlighting its ecological adaptability across several hosts [25–27]. The recently described environmental species *E. aeris* can solubilize organic phosphate and produce siderophore [28]. Although primarily known as plant- or insect-associated bacteria, members of the genus have also been found associated with rare human infections; for example, *E. billingiae* has been reported as a causal agent of septic arthritis [29]. *E. billingiae* strains are more commonly isolated from stem cankers, diseased blossoms, and immature fruits of rosaceous trees, where they typically act as secondary invaders rather than primary pathogens [30,31].

*Erwinia* species possess a diverse array of virulence determinants that enable host colonization, modulation of host defenses, and disease progression. These factors include different type secretion systems, exopolysaccharide biosynthetic pathways, iron-uptake systems, flagella-mediated motility, and diverse gene clusters that contribute to pathogenicity and ecological adaptation [7, 32–34]. Previous studies indicate that several virulence determinants in the genus *Erwinia*, including levan biosynthesis, sorbitol metabolism, and multiple type III and type VI secretion systems, have an ancestral origin, whereas other factors, such as a second flagellar system and glycosyltransferases involved in amylovoran biosynthesis, were likely acquired later during the evolution of pathogenic lineages [35]. However, understanding the evolutionary distribution and diversification of these determinants requires comprehensive comparative genomic and phylogenomic analyses [36]. Advances in high-throughput sequencing have greatly expanded the number of available *Erwinia* genomes, enabling robust genome-scale analyses based on core genes. Such genome-wide approaches improve taxonomic resolution, facilitate species delineation, and provide insights into the evolutionary processes shaping genomic diversity across the genus [37].

Despite the increasing availability of genomic data, comparative studies of the genus *Erwinia* remain limited in scope and have primarily focused on a few pathogenic species [38–42]. To our knowledge, no study has comprehensively integrated genus-wide phylogenomics, species delineation, comparative analyses of virulence factors, antimicrobial gene clusters, and other genome-scale features across the genus. To address this knowledge gap, the present work provides the comprehensive phylogenomic and comparative genomic analyses of the genus *Erwinia*. By incorporating representative genomes from all currently recognized species, we present the evolutionary relationships across the genus and assess key genomic features associated with both pathogenic and non-pathogenic lifestyles. Also, we identified a genomically distinct lineage isolated from dogwood (*Cornus florida*) that falls below established species delineation thresholds based on average nucleotide identity (ANI) and digital DNA-DNA hybridization (dDDH), suggesting that it may represent a previously undescribed lineage within the genus *Erwinia*, potentially corresponding to a new species.

## Material and Methods

### Bacterial strains, culture conditions and genomic DNA extraction

A strain of *Erwinia* sp. PL328 originally isolated from dogwood (*Cornus florida*) was obtained from the Pacific Bacterial Culture Collection (University of Hawai’i at Manoa, Honolulu, HI, United States). The genomic DNA was isolated using the DNeasy Blood and Tissue Kit (Qiagen, Germantown, MA) following the manufacturer’s instructions. Strain identification was performed by amplification of 16SrRNA gene region using the universal primers P16S F and P16S R under the PCR conditions as described by Larrea-Sarmiento et al. [43]. The amplified product was sequenced by Sanger sequencing at the GENEWIZ facility (Azenta Life Sciences, La Jolla, CA), and the resulting sequence was queried against the NCBI nucleotide database using BLASTn, confirming its identity as *Erwinia* sp. One strain of *Erwinia amylovora* (LMG 1877) was obtained from the BCCM-LMG culture collection (Ghent, Belgium). This strain was selected because it represents one of the earliest available *E. amylovora* isolates, collected in 1972 from *Cydonia oblonga*, and because its genome sequence was not available in the NCBI database. Therefore, LMG 1877 served as a valuable historical reference strain for the study, particularly against the type strain *E. amylovora* ATCC 15580^T^.

The dogwood strain PL328 was cultured on tetrazolium chloride agar (TZC: Peptone 10 g/L, dextrose 5 g/L, agar 17 g/L and 0.001% TZC) [44], while *Erwinia amylovora* LMG 1877 was cultured on Medium 6 (Glucose 10g/L, Yeast extract 5g/L, Peptone 5g/L, Agar 15g/L, pH adjusted to 7.0) as recommended by the BCCM/LMG Bacteria Collection (Belgian Co-ordinated Collections of Micro-organisms, Laboratory of Microbiology, Department of Biochemistry and Microbiology, Faculty of Sciences of Ghent University), and both were incubated at 28 °C for 24 h. To ensure culture purity, a single colony was restreaked and incubated under the identical conditions. Whole genomic DNA of these two strains was extracted from approximately half a loopful of overnight-grown culture using the QIAGEN Genomic-tip 100/G kit (Qiagen, Valencia, CA) following the manufacturer’s instructions. DNA quality and concentration were assessed using a Nanodrop spectrophotometer and a Qubit 4 fluorometer (Thermo Fisher Scientific, Life Technologies, Carlsbad, CA).

In addition to these two strains sequenced in this study, 102 *Erwinia* genomes representing the genomes from different species were retrieved from the NCBI GenBank genome database. These genomes originated from approximately 25 geographically diverse regions, including China, United States, Italy, Germany, United Kingdom, Portugal, South Korea, Canada, France, Poland, New Zealand, Japan, Denmark, Brazil, and Australia, and were isolated from approximately 42 different host species and ecological niches, including *Malus domestica*, *Cydonia oblonga*, *Crataegus* sp., *Allium cepa*, *Pyrus communis*, *Carica papaya*, *Brassica rapa*, *Solanum tuberosum*, *Cucumis sativus*, soil and environment **(Supplementary Table 1)**.

### Whole-Genome Sequencing and Annotation

The whole-genome sequencing of *Erwinia* sp. dogwood isolate PL328 and *E. amylovora* LMG1877 strains was performed by Plasmidsaurus (San Francisco, CA) using Oxford Nanopore long-read and Illumina short-read sequencing technologies. For long-read sequencing, libraries were prepared using the Rapid Barcoding Kit 96 V14 and sequenced on a PromethION P24 platform with R10.4.1 flow cell. Basecalling was conducted using Dorado v7.1.4 [45]. Reads with quality score below 10 were removed and adapter sequences were trimmed using MinKnow. *De novo* assembly was performed using Flye v2.9.1 [46] and subsequently polished with Medaka v1.8.0 [47].

Briefly, for Illumina short-read sequencing, libraries were prepared using the Illumina DNA Prep Kit and sequenced on a NextSeq 2000 platform with paired-end 2 × 150 bp reads. The Oxford Nanopore assemblies were polished using Illumina short-reads with Polypolish v0.6.0 [48]. The final assemblies were annotated using Bakta v1.6.1 [49], the NCBI Prokaryotic Genome Annotation Pipeline (PGAP v6.10) [50], and the Bacterial and Viral Bioinformatics Resource Center (BV-BRC v3.54.6) [51], and the assembled genomes were subsequently submitted to the NCBI GenBank database.

### Genome similarity and taxonomic resolution

The genome sequences of the type strain of all species and the two strains sequenced in this study were included in this analysis, bringing the total to 26 strains. When the genome of a type strain was unavailable, the closest representative genome was selected from the NCBI GenBank database (accessed June 10, 2025) **(Supplementary Table 2)**. Pairwise comparisons between LMG 1877, the dogwood isolate PL328, and the other 24 genomes were performed using average nucleotide identity (ANI) calculated with the FastANI algorithm v1.34 [52]. Species boundaries were further confirmed using digital DNA–DNA hybridization (dDDH) estimated through the Type Strain Genome Server (TYGS) [53]. Species delineation was interpreted using the generally accepted thresholds of 95% ANI and 70% dDDH [54–59]. The ANI and dDDH values were compiled into a single matrix and visualized as a color-coded heatmap using Displayr (https://www.displayr.com).

### Phylogenetic analysis

A total of 104 complete and high-quality draft genome sequences representing the currently available genomic diversity of the genus *Erwinia* were included in the analyses **(Supplementary Table 1)**. All the genome sequences were re-annotated using Prokka [60], and the resulting GFF files were used for downstream pan-genome analyses. The pan- and core-genome analyses were performed using Roary v3.13.0 [61]. Preliminary attempts to run Roary with higher BLASTp identity thresholds (90-95%) consistently failed to generate a core genome alignment, likely due to extensive genomic heterogeneity across *Erwinia* species, therefore a cutoff of 85% was used. It was run with 85% BLASTp identity and 90% core threshold, and the core gene sequences were aligned with MAFFT (Multiple Alignment using Fast Fourier Transform) for the construction phylogenomic tree. Genes present in ≥90% of genomes were classified as core genes, whereas genes present in ≥89% and <90%, ≥15% and <89%, and <15% of genomes were classified as soft-core, shell, and cloud genes, respectively. A pan-core genome plot of 104 genomes was generated using ggplot in R, based on the roary output to visualize pan and core-genome dynamics across the dataset. The resulting core-genome alignment was used for phylogenetic reconstruction. A maximum-likelihood phylogenetic tree was constructed using IQ-TREE2 v2.1.2 [62]. The best-fitting nucleotide substitution model was selected using ModelFinder according to the Bayesian Information Criterion (BIC). The GTR+I+G model was identified as the optimal model and used for tree reconstruction. Branch support was assessed using 1,000 ultrafast bootstrap replicates. The final phylogenetic tree was visualized and annotated using iTOL v7 [63].

### Comparative genomics

Comparative genomic analyses were performed using 26 strains representing type/reference strains representing the type or reference strains of recognized *Erwinia* species together with the genomes sequenced in this study **(Supplementary Table 2)**. For species-level pan-genome and orthologous gene analyses, a non-redundant dataset comprising 21 representative genomes (one representative strain per species, preferably the type strain or an appropriate reference genome when the type-strain genome was unavailable) was used. The pan and core genome analyses were performed using Roary v3.13.0 [61] with a BLASTp identity threshold of 80% [64]. The strict core, accessory, strain-specific, and total gene counts for each strain were derived from the Roary output. The strict core genome comprised genes present in all 21 strains, whereas strain-specific genes were present exclusively in a single strain. The accessory genome comprised genes present in two or more strains, but not in all 21 strains. Flower plots illustrating core, accessory, strain-specific and total gene distributions were generated using custom scripts in R.

To investigate the evolution of genes across different species, orthologous gene clusters were identified among the same 21 representative genomes using the OrthoVenn 3 [65], which uses the OrthoMCL algorithm with a default E-value of 1 x 10^-2^. Genomes were analyzed into sets of four or five based on their phylogenetic proximity as inferred from the phylogenetic tree. The focal clade consisting of *E. amylovora*, *E. pyrifoliae*, *E. piriflorinigrans*, and *E. tasmaniensis* was analyzed independently and together with the adjacent sister species *E. aphidicola* to assess orthologous gene conservation within the focal clade and in the context of its closest evolutionary relative. To characterize genome architecture and horizontally acquired regions, genomic islands (GIs) were predicted for all genomes included in **Supplementary Table 2** using IslandViewer4 with default parameters [66]. IslandViewer4 integrates multiple approaches, including IslandPick (comparative genomics), IslandPath-DIMOB and SIGI-HMM (sequence composition-based methods), and Islander (site-specific integration into tRNA/tmRNA genes) [66]. The genome annotations for the strains were performed using Bacterial and Viral Bioinformatics Research Center (BV-BRC) [51], and the resulting GenBank files were used as an input for GI prediction in the strains.

Prophage-like regions were identified in the bacterial genomes, listed in **Supplementary Table 2**, using the Phage Search Tool Enhanced Release (PHASTER; University of Alberta, Canada) [67,68]. Prophages are important contributors to genomic diversification and may influence differences in virulence and host adaptation across species [69]. The antimicrobial resistance genes were identified using RESFinder v 4.7.2, which uses BLAST algorithm to compare the genome sequences against a curated database of antimicrobial resistant genes [70]. This analysis was performed to evaluate the distribution of resistance determinants across *Erwinia* species.

For the 26 genomes listed in **Supplementary Table 2,** pathogenicity-associated gene clusters involved in type III and type VI secretion systems, amylovoran biosynthesis, cellulose production, levan metabolism, sorbitol metabolism, iron-scavenging siderophores (desferrioxamine and proferrorosamine), metalloproteases and flagellar assembly were retrieved using the locus tags reported in the study conducted by Smits et al. [71], Smits et al. [72] and Kube et al. [73]. These clusters were compared among the *Erwinia* species using the Geneious Prime v 2025.2 (https://www.geneious.com). Synteny and genomic rearrangements among major pathogenicity-associated gene clusters were visualized using the clinker tool implemented in the Comparative Gene Cluster Analysis Toolbox (CAGECAT) using its clinker tool [74]. To identify pathogenicity-associated genes and clusters that could not be recovered through annotation-based extraction, one-versus-one BLASTp analyses were performed using the complete genome of *E. amylovora* ATCC 15580^T^ as the reference, allowing determination of the percent identity across all *Erwinia* strain genomes in this study [75,76]. For the proferrorosamine biosynthesis (r*osA*-*rosG*) cluster, protein sequences from *E. rhapontici* P45 were used as the reference queries [5]. An e-value threshold of 1e^-2^ was used to ensure retention of genuine but divergent homologs, preventing false-negative loss of variably conserved pathogenicity genes across *Erwinia* species [77].

## Results

### Genomic and phylogenetic analyses

The complete genomes of the dogwood isolate PL328 and *E. amylovora* LMG1877 consisted of a chromosome of size 4.92- and 3.80-Mb, respectively, and both exhibited a GC content of 53.5%. Genome annotation predicted 4,954 coding sequences together with 22 rRNA and 84 tRNA genes in PL328 and 3,923 coding sequences, 22 rRNA and 77 tRNA genes in LMG1877 genome. Two CRISPR arrays were identified in LMG1877, whereas no CRISPR arrays were detected in the dogwood isolate PL328. Both assemblies showed high completeness (>99% coarse consistency; 100% CheckM completeness), supporting their suitability for downstream comparative genomic analyses.

The genomic relatedness among the *Erwinia* genomes based on 26 strains (**Supplementary Table 2)** revealed substantial genetic diversity within the genus with ANI and dDDH ranging between 79.0 to 100% and 20.2 to 99.9%, respectively (**Figure 2**). The *Erwinia* sp. PL328 isolated from dogwood exhibited ANI and dDDH values below the accepted species delineation thresholds (<95% ANI and <70% dDDH) [54–59] relative to all known *Erwinia* species, indicating a potential new species. *E. aeris* ACC02193^T^ formed a group with *E. plantamica* OPT-41^T^ and *E. persicina* ZSR3, sharing 99.1% ANI and 93.9-94% dDDH values, indicating that these strains belong to the same genomic species. Similarly, *E. papayae* JGD 233 grouped with *E. mallotivora* ICMP5705^T^ and shared 96.9% ANI and 75.2% dDDH, consistent with their placement within a single genomic species. However, *E. persicina* strains CFBP 8797 and CFBP 8803 exhibited ANI and dDDH values below the accepted species delineation thresholds when compared with the type strain *E. persicina* NBRC 102418^T^ and all other *Erwinia* species. Similarly, *E. tasmaniensis* PPS120 showed ANI and dDDH values below the species-level thresholds compared with type strain *E. tasmaniensis* Et1/99^T^ and remaining *Erwinia* species, suggesting that these strains may represent potential novel species within the *Erwinia* genus. A maximum-likelihood phylogenetic tree based on the core genome was constructed using IQ-TREE2 v2.1.2 [62] under the GTR + I + G model with 1,000 ultrafast bootstrap (**Figure 3**).

**Figure 2.**
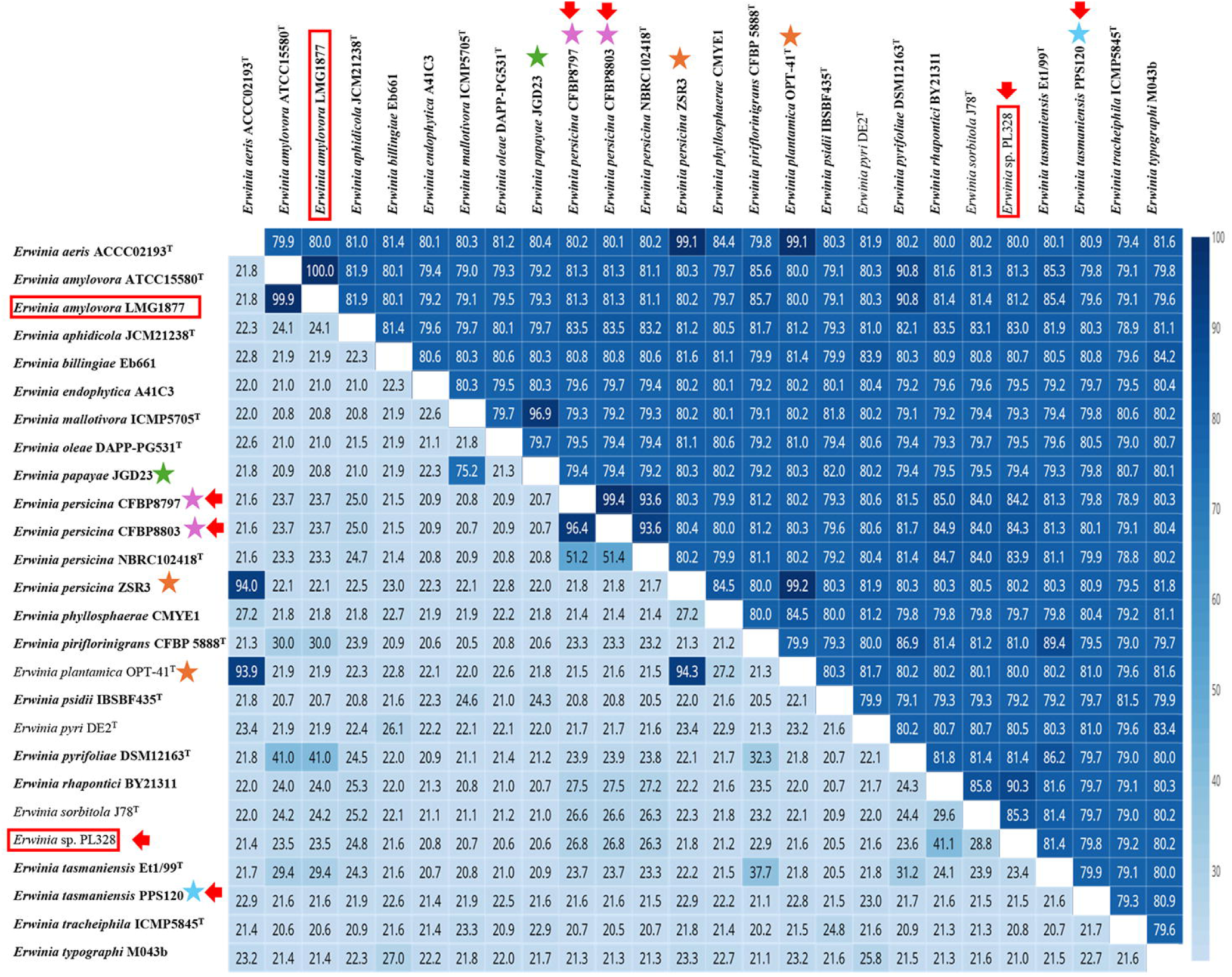
Heatmap of pairwise genomic relatedness across the genus measured by Average Nucleotide Identity (ANI, top triangle) and digital DNA–DNA hybridization (dDDH, bottom triangle). Color gradient ranges from light (low percent similarity) to dark (high percent similarity). Values inside cells are percent ANI or percent dDDH. Colored star symbols (★) indicate potentially misclassified strains; each color represents a distinct misclassification group. Boxed strains represent genomes reported in this study, and the red colored arrow represents the new *sp*. reported from this study.

**Figure 3.**
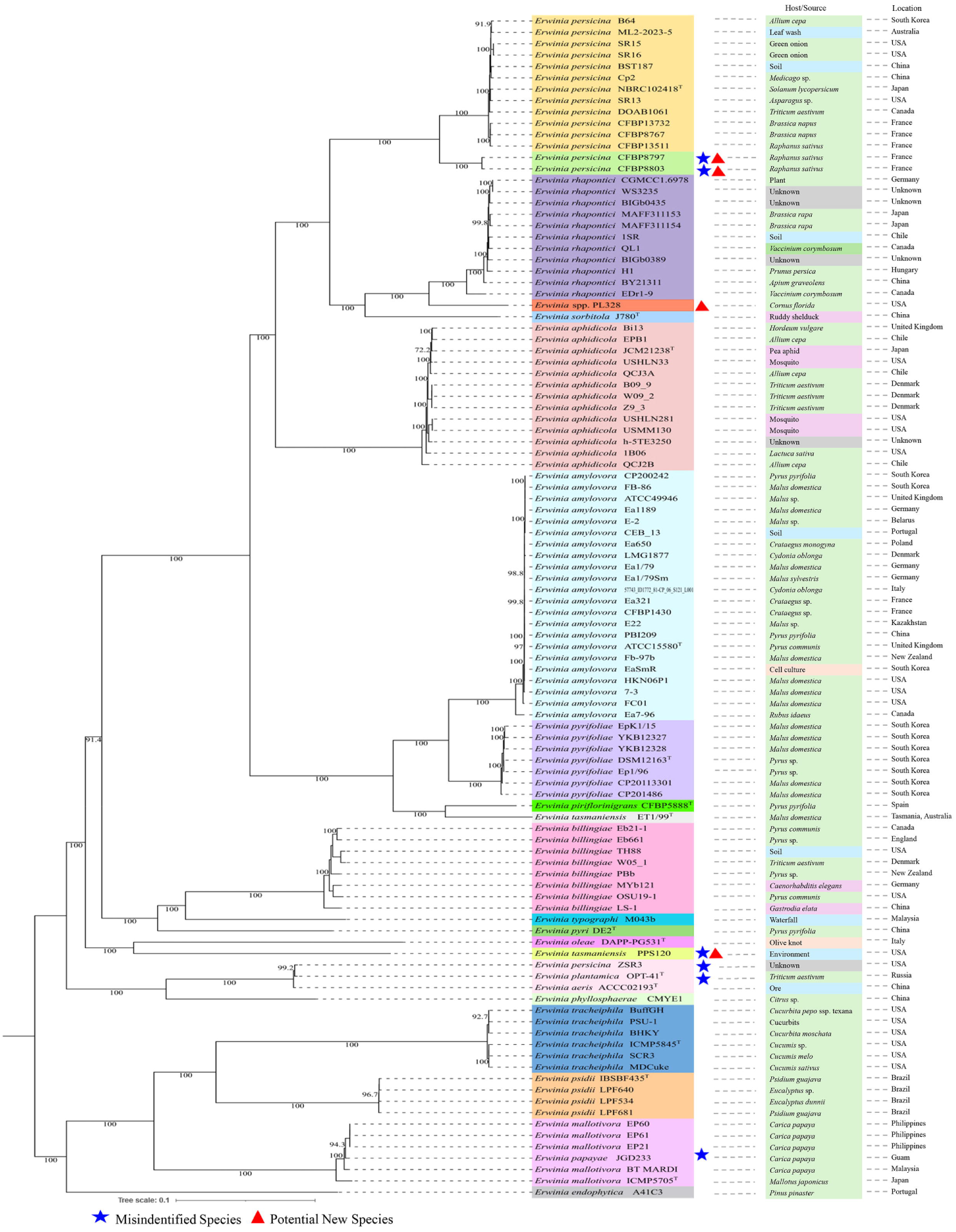
Maximum likelihood phylogenetic tree showing relationships among *Erwinia* strains generated using IQTree2 v2.1.2 and visualized using iTOL annotation and editor v7. Stars indicate putatively misidentified taxa, whereas triangles denote potential novel species identified in this study. Two annotation boxes are shown adjacent to each terminal taxon: the left box represents the host of isolation, and the right box represents the geographic origin of the isolate. Host/source categories are color-coded as follows: plant (green), environment (blue), insect (pink), other hosts/source (orange), and unknown (gray).

The phylogenomic analysis based on core-genome alignment clearly resolved species-level relationships within the genus *Erwinia*, with high bootstrap support values across major clades, indicating reliable evolutionary separation, and also confirming the ANI and dDDH based species delineation. The analysis revealed a distinct and well-supported clade formed by the strains of *E. persicina*. However, two strains (*E. persicina* CFBP 8797 and CFBP 8803) formed a separate sub-cluster, presenting some distinctness in their genome sequences. The ANI and dDDH values of these strains also suggest that they may represent a novel species within the *Erwinia* genus. Adjacent to this group, *E. rhapontici* isolates formed a highly supported monophyletic cluster, reflecting low genomic variability within this lineage. The dogwood isolate, ‘*Erwinia* sp. PL328’, formed a distinct clade, indicating a unique evolutionary position and suggesting it may represent a novel *Erwinia* lineage. The *E. amylovora* strains grouped into a monophyletic cluster with high intra-species similarity (ANI > 99%, dDDH > 70%), while *E. pyrifoliae* strains, *E. piriflorinigrans* CFBP 5888^T^ and *E. tasmaniensis* Et1/99^T^ appeared as neighboring but distinct branches. Interestingly, the *E. tasmaniensis* PPS120 branched independently, clustering near *E. oleae* DAPP-PG531^T^, consistent with the ANI and dDDH analyses and suggesting that *E. tasmaniensis* PPS120 represents a distinct genomic lineage from *E. tasmaniensis* Et1/99^T^. Also, *E. persicina* strain ZSR3 and *E. plantamica* strain OPT-41^T^ grouped with *E. aeris* ACC02193^T^, suggesting either misclassification in their taxonomic placement and representing a common lineage within the *E. aeris* group. Additionally, *E. papayae* JGD 233 clustered closely with *E. mallotivora* ICMP5705^T^ with strong bootstrap support, indicating a close evolutionary relationship between these pathogens and suggesting that *E. papayae* JGD 233 may require reassessment or potential reclassification within the *E. mallotivora* lineage.

Overall, the core-genome phylogeny, together with ANI and dDDH analyses confirmed recognized species boundaries as resolved in this study (**Figure 2 and 3**) within the genus. The distinct placement of the dogwood isolate PL328, *E. persicina* strains CFBP 8797 and CFBP 8803, and *E. tasmaniensis* PPS120 suggests the presence of previously unrecognized species-level lineages. In addition, the clustering of *E. persicina* ZSR3 and *E. plantamica* OPT-41^T^ with *E. aeris* ACC02193^T^ indicates a shared genomic lineage and possible misclassification, while the close relationship between *E. papayae* JGD 233 and *E. mallotivora* ICMP5705^T^ supports the need for reassessment of their current taxonomic status. Collectively, these findings highlight the complexity of species boundaries within the genus *Erwinia* and underscore the need for further taxonomic refinement.

### Pan-core genome analysis of the genus *Erwinia*

To complement the phylogenomic analysis and further explore genomic diversity within the genus *Erwinia*, a pan-core genome analysis was conducted using 104 *Erwinia* genomes and the resulting conserved and total gene distribution is presented in **Supplementary Figure 1**. The pan-genome comprised a total of 56,424 gene families, of these, the core genome, defined as genes present in ≥90% of strains consisted of 1,036 gene families. An additional 14 soft-core genes were identified, present in 89-90% of genomes. In contrast, 4,741 genes were classified as shell genes, and a large proportion (50,633 genes) formed the cloud genome, being present in fewer than 15% of strains.

To study the species level distinctness, this analysis was further performed using only the type or representative strains from each species (21 strains). This subset revealed a strict core genome of 1,174 genes, while the total and strain-specific gene counts varied substantially among the species (**Figure 4**). *E. tracheiphila* ICMP5845^T^ possessed both the highest total gene count and the largest number of strain-specific genes. The unique gene repertoire included cell-wall degrading *cbhA* and *celZ*, pectin-degrading rhamnogalacturonate lyase, *rhiE* [78,79], the expansin *YoaJ*, a virulence-associated protein previously demonstrated to enhance bacterial wilt development in *E. tracheiphila* [80], together with numerous genes representing diverse predicted functions, including several proteins of unknown function, followed by *E. typographi* M043b. These counts may reflect lineage-specific gene content variation, although assembly fragmentation and annotation-related effects associated with draft genomes may also contribute to high unique gene estimates. In contrast, the highest number of accessory genes was observed in *E. persicina* NBRC 102418 ^T^ (2,911 genes), followed closely by *E. persicina* CFBP 8797 (2,888 genes). Across nearly all species (except *E. tracheiphila* ICMP5845^T^ and *E. oleae* DAPP-PG531^T^), the accessory genome exceeded the size of the core and unique gene fractions, indicating substantial accessory genome diversity within the genus *Erwinia*.

**Figure 4.**
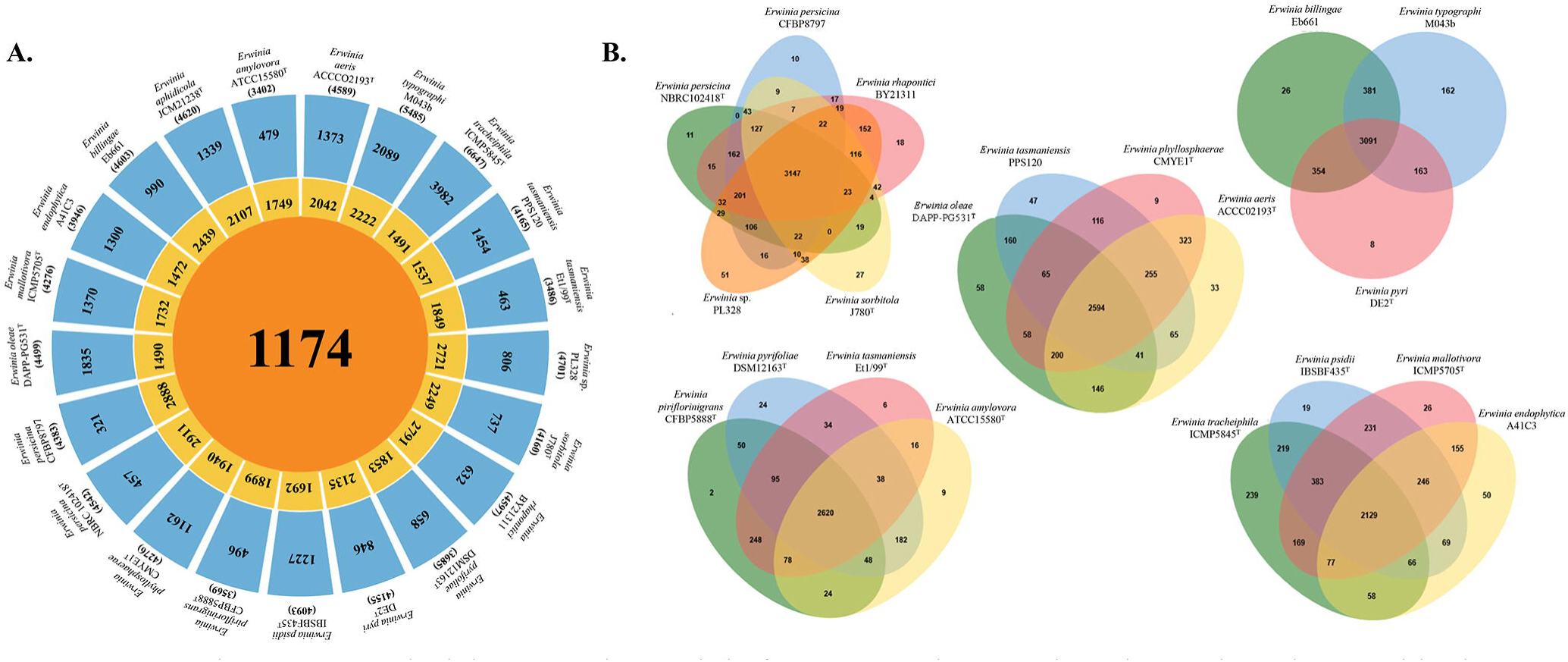
Comparative pan-genome and orthologous gene cluster analysis of *Erwinia* type and representative strains. **(A)** Flower plot summarizing the pan-core genome of *Erwinia* type and representative strains. For each strain, the center indicates the number of core genes, the annulus represents accessory genes, and the petal tips denote unique genes. The total number of genes for each strain is shown outside the petals next to the strain name. **(B)** Orthologous gene cluster comparison among *Erwinia* type and representative strains using OrthoVenn3. Strains were grouped according to their phylogenetic proximity, and orthologous clusters were identified using the OrthoMCL algorithm. Each multi-set Venn diagram displays the distribution of shared and group-specific orthologous gene clusters within a phylogenetic group. Overlapping regions represent clusters conserved across multiple strains, while non-overlapping sections correspond to lineage-specific orthologous clusters.

### Comparative genomics

Orthologous gene clusters were analyzed among the 21 representative *Erwinia* genomes (one representative strain per species) to identify conserved and lineage-specific genomic content across the genus (**Figure 4**). Across the genus, a substantial set of conserved orthologs formed the essential metabolic and functional backbone of the genus *Erwinia*, while species grouped within the same phylogenetic clades shared a greater proportion of orthologous genes, consistent with their close evolutionary relationships and similar ecological adaptations. The clade including *E. persicina* NBRC102418^T^ and *E. persicina* CFBP8797, *E. rhapontici* BY21311, *E. sorbitola* J780^T^, and the *Erwinia* sp. PL328 isolated from dogwood in this study exhibited the largest shared core gene set (3,147 genes) with relatively few species-specific genes, indicating strong genomic conservation. Notably, the *Erwinia* sp. PL328 isolated from dogwood possessed the highest number of species-specific genes (51 genes) within this clade, including genes associated with amino acid transport, DNA topology, DNA binding proteins, and numerous proteins of unknown function, supporting its genomic distinctiveness and a potential novel lineage.

Among the pathogens isolated from pome fruit (*E. amylovora* ATCC15580^T^, *E. pyrifoliae* DSM12163^T^, *E. piriflorinigrans* CFBP5888^T^ and *E. tasmaniensis* Et1/99^T^) as shown in **Figure 4**, a high overlap in gene content was observed (2,620 orthologous genes), predominantly representing functions associated with transport, transcriptional regulation, protein secretion (including the type III secretion system), pathogenesis, motility, and core metabolism, with only 9, 24, 2 and 6 species-specific genes, respectively. When the insect-associated species *E. aphidicola* JCM21238^T^ placed in a neighbouring sister clade was included in the analysis **(Supplementary Figure 2)**, the number of orthologous genes reduced to 2,456, while *E. aphidicola* JCM21238^T^ contained the maximum number of strain-specific genes (183 genes) in this group. The *E. billingiae* Eb661-*E. typographi* MO43b-*E. pyri* DE2^T^ group shared a core of 3,091 proteins, but showed considerable variation in species-specific genes (26, 162 and 8 genes) respectively. The substantially higher number of species-specific genes in *E. typographi* MO43b was associated with mobile genetic elements, phage-related functions and stress responses. The lineage comprising *E. psidii* IBSBF435^T^, *E. tracheiphila* ICMP5845^T^, *E. mallotivora* ICMP5705^T^, and *E. endophytica* A41C3 contained the smallest core genome (2,129 genes) together with a high accessory gene count, indicative of broad ecological diversification, including plant pathogens and endophytes. The expanded species-specific gene repertoire of *E. tracheiphila* ICMP5845^T^ was dominated by mobile genetic element- and bacteriophage-associated functions, whereas those of *E. endophytica* A41C3 were associated with iron acquisition, nutrient transport, cell adhesion, and diverse metabolic functions. The non-plant pathogenic cluster (*E. oleae* DAPP-PG531^T^, *E. tasmaniensis* PPS120, *E. phyllosphaerae* CMYE1^T^, and *E. aeris* ACCC02193^T^), shared a clade-specific set of 2,594 genes and possessed very few unique genes (9-58 genes), primarily associated with DNA mobility, nutrient transport, stress response, iron acquisition, transcriptional regulation, and central metabolism. Collectively, the ortholog comparison demonstrate the presence of a conserved core genome across *Erwinia*, while the presence of lineage-specific gene pools suggests evolutionary diversification linked to host range, ecology, and niche specialization across the genus [81,82].

Genomic island (GI) prediction across all 26 *Erwinia* genomes revealed notable variation in horizontally acquired regions across the *Erwinia* genomes. *Erwinia pyrifoliae* DSM12163^T^ possessed the highest number of GIs (56 islands), followed by *E. typographi* MO43b (52 islands), whereas *E. pyri* DE2^T^ and *E. phyllosphaerae* CMYE1^T^, and contained the least 17 and 21 islands, respectively presented in **Supplementary Table 3**. Among the genomes reported in this study, the potential novel *Erwinia* sp. isolate carried 35 GIs and *E. amylovora* LMG1877 exhibited 23 islands, reflecting comparatively moderate levels of genomic island acquisition.

### Prophage and Phage-like elements within the *Erwinia* genus

PHASTER analysis revealed the presence of intact prophages and prophage-like elements across all 26 analyzed *Erwinia* genomes. At least one putative complete (intact) prophage was detected in nearly all species, with the exception of *E. amylovora* ATCC 15580^T^, *E. amylovora* LMG 1877, *E. phyllosphaerae* CMYE1^T^, *E. sorbitola* J780^T^, and *E. tasmaniensis* Et1/99 ^T^, which lacked intact prophages but have incomplete and questionable elements. *E. amylovora* LMG 1877 contained three incomplete prophages measuring approximately 8.5 kb, 27.6 kb, and 23.2 kb. In contrast, *Erwinia* sp. PL328 isolated from dogwood possessed one intact prophage (16.3 kb) with 23 predicted proteins), along with three incomplete prophages. Interestingly, bacterial-origin genes within prophage regions were detected only in two *Erwinia* species, including *E. billingae* Eb661 and *E. endophytica* A41C3 whereas the remaining genomes lacked such features (**Supplementary Table 4**). This suggests variability in prophage acquisition and retention among *Erwinia* species, suggesting differential prophage-associated genomic features across the genus. Antimicrobial resistance determinants were identified only in *E. persicina* NBRC102418^T^ and *E. persicina* CFBP8797 specifically against amoxicillin, ampicillin, piperacillin and ticarcillin **(Supplementary Table 5)**.

### Type secretion system and other virulence factors

Comparative genomic analysis of all 26 *Erwinia* genomes identified multiple virulence-associated determinants across the *Erwinia* genus, including type III and type VI secretion systems, extracellular polysaccharide biosynthesis clusters (amylovoran, levan, and cellulose), metalloproteases, flagellar gene clusters, sorbitol metabolism genes, and iron acquisition systems (**Supplementary Table 6**). One-versus-one BLASTp analyses [75,76] revealed variation in the distribution of pathogenicity-associated genes and gene clusters across the pathogenic and non-pathogenic *Erwinia* species evaluated in this study (**Supplementary Table 6**).

Three Type III secretion system (T3SS) clusters were identified, including Pathogenicity Island 1 (PAI-1), comprising the canonical Hrp-T3SS (*hrp*/*hrc*, *hee*, and *hae* regions), and two Inv/Spa-type T3SS clusters (Pathogenicity Island II and III, respectively). Comparative analysis revealed a conserved *hrp*-T3SS (PAI-1) architecture centered on the canonical *hrp/hrc* gene cluster, in *E. amylovora* ATCC15580^T^ and LMG1877, *E. pyrifoliae* DSM12163^T^, and *E. tasmaniensis* Et1/99^T^, except for *orfU1* and YgjP-like metallopeptidase domain-containing protein, which were unique to *E. amylovora* ATCC15580^T^ and LMG1877. The *hrp/hrc* region is flanked by the *hae* and *hee* regions, which harbor additional virulence-associated genes. *E. tasmaniensis* Et1/99^T^ lacked the *hae* region, while three of the species (*E. amylovora* ATCC15580^T^ and LMG1877, *E. pyrifoliae* DSM12163^T^, and *E. tasmaniensis* Et1/99^T^) retained a complete *hee* region (**Figure 5A**). The distribution of Inv/Spa-type T3SS islands (PAI-2 and PAI-3) was more variable across *Erwinia* species. Both the islands were present in *E. amylovora* ATCC15580^T^ and LMG1877, whereas the PAI-2 T3SS was absent in *E. pyrifoliae* DSM12163^T^ and PAI-3 was partially lost in *E. tasmaniensis* Et1/99^T^, indicating independent loss events after species divergence (**Figure 5A**). The blastp analysis revealed an almost complete PAI-1 and PAI-2 regions in *E. piriflorinigrans* CFBP 5888^T^ with > 79% identity and > 65% query coverage relative to *E. amylovora* ATCC15580^T^, suggesting lineage-specific retention linked to closely related species of *Erwinia* [35,83]. In contrast, only homologs of few T3SS-associated proteins were detected in *E. mallotivora* ICMP5705^T^ (74–84% identity), *E. oleae* (77–88%), *E. phyllosphaerae* CMYE1 (75–79%), *E. psidii* IBSBF435^T^ (73–83%), and *E. tracheiphila* ICMP5845^T^ (78–83%) (**Supplementary Table 6)**. Complete PAI-1 and Inv/Spa-type T3SS loci could not be reconstructed in these species using *E. amylovora* ATCC15580^T^ as the reference. Additionally, three effector singletons (*eop2*, *hopPtoC*, and *avrRpt2*), potentially contributing to increased virulence were found exclusively in *E. amylovora* ATCC15580^T^ and LMG1877.

**Figure 5.**
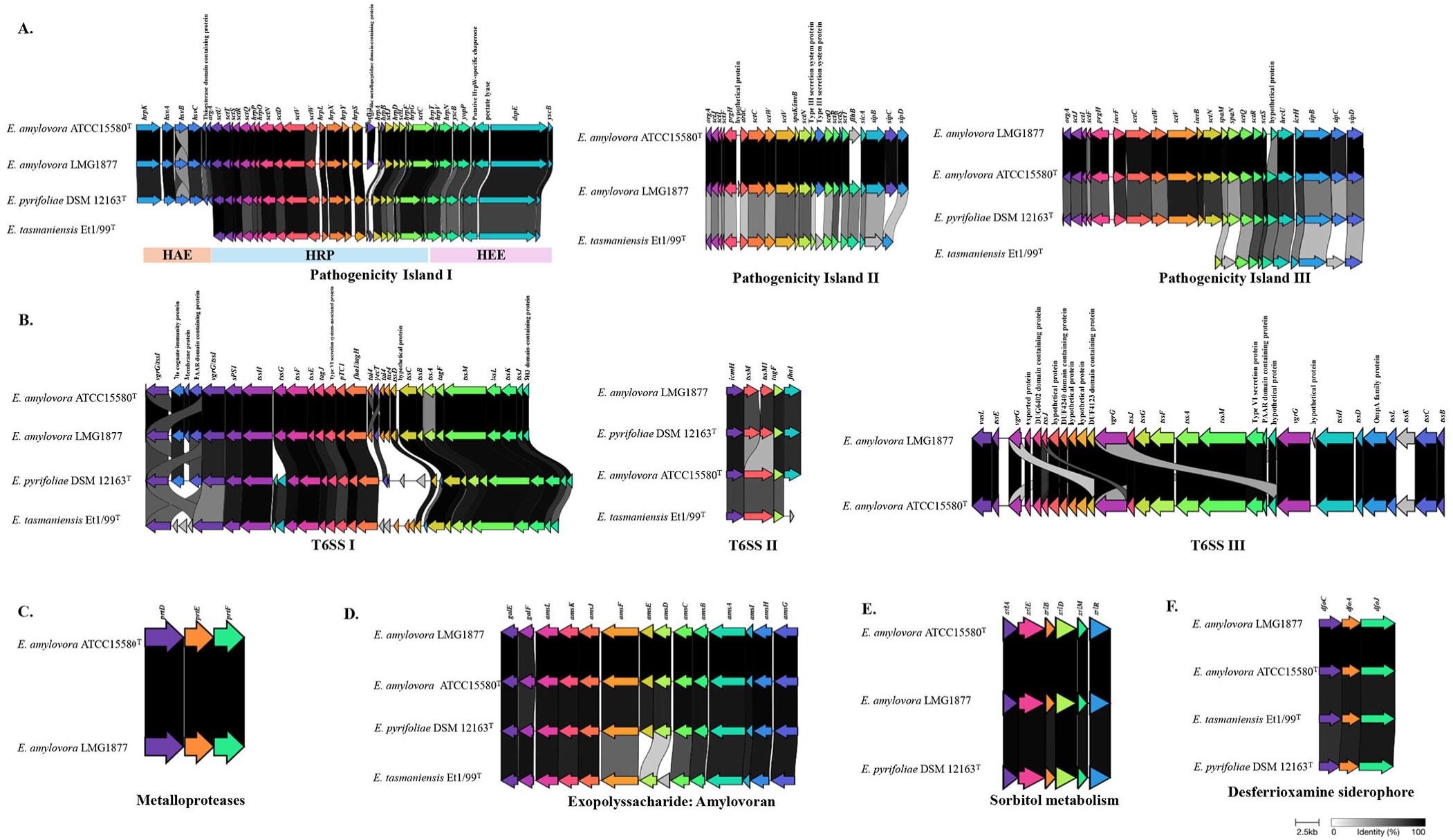
Distribution of secretion system-associated gene clusters among *Erwinia* species identified by reference mapping in Geneious Prime v 2025.2. (A) Distribution and comparative organization of Type III secretion system (T3SS) gene clusters among different *Erwinia* species (B) Distribution and comparative organization of Type VI secretion system (T6SS) gene clusters among different *Erwinia* species (C) Sorbitol metabolism-associated gene cluster among different *Erwinia* species (D) Exopolysaccharide biosynthesis (amylovoran) gene cluster among different *Erwinia* species (E) Desferrioxamine siderophore biosynthesis-associated gene cluster among different *Erwinia* species (F) Metalloprotease-associated gene cluster among different *Erwinia* species.

Three distinct Type VI secretion system (T6SS) clusters were identified in the genus *Erwinia*. The T6SS-1 cluster represents a complete structural system comprising all core components (*tssA-M*), *tagF/H/J* together with multiple effector-immunity pairs (*Tae4/Tai4*). T6SS-2 appears to be a degenerate or partially duplicated cluster derived from the ancestral T6SS-1 locus (De Maayer et al., 2011). The analyses revealed that T6SS-1 and T6SS-2 are conserved among *E. amylovora* ATCC15580^T^ and LMG1877, *E. pyrifoliae* DSM12163^T^, and *E. tasmaniensis* Et1/99^T^ (**Figure 5B**). Outside this group, homologs of several core T6SS-1 and T6SS-2 structural components were detected in multiple *Erwinia* species using blastp analysis **(Supplementary Table 6)**. However, complete T6SS loci could not be mapped using *E. amylovora* as a reference maybe due to variable conservation and differences in locus organization among species. The T6SS-3 was detected exclusively in *E. amylovora* ATCC15580^T^ and LMG1877 (**Figure 5B**). The PrtDEF Type I secretion system (T1SS) encodes secreted metalloprotease PrtA and its export apparatus (PrtD, PrtE, PrtF). This cluster was identified exclusively in the genome of *E. amylovora* ATCC15580^T^ and LMG1877 and was absent from all other *Erwinia* species analyzed (**Figure 5C**).

Also, the exopolysaccharide amylovoran biosynthetic cluster, composed of 14 *ams* genes, is conserved in *E. amylovora* ATCC15580^T^ and LMG1877, *E. pyrifoliae* DSM12163^T^ and *E. tasmaniensis* Et1/99^T^. In *E. pyrifoliae* DSM12163^T^, the exopolysaccharide is encoded by the cps operon, which is orthologous to the *ams* operon in *E. amylovora* ATCC15580^T^ and LMG1877, while *E. piriflorinigrans* CFBP5888^T^possessed a conserved locus lacking the *amsD* gene (**Figure 5D**). The BLASTp analysis identified homologs of several *ams* proteins in multiple *Erwinia* species **(Supplementary Table 6).** However, complete amylovoran biosynthetic clusters were not identified except in the species described above (*E. amylovora* ATCC15580^T^ and LMG1877, *E. pyrifoliae* DSM12163^T^, *E. tasmaniensis* Et1/99^T^ and *E. piriflorinigrans* CFBP5888^T^). No homologs of the *ams* genes were detected in *E. persicina* ZSR3, *E. aeris* ACCC02193^T^, and *E. plantamica* OPT-41. The cellulose biosynthesis operon (*bcsA*-*bcsD*) was fully conserved in *E. amylovora* ATCC15580^T^ and LMG1877, and *E. pyrifoliae* DSM12163^T^, whereas one gene *(bcsD)* was missing in *E. tasmaniensis* Et1/99^T^, *E. piriflorinigrans* CFBP5888^T^, *E. aphidicola* JCM21238^T^, *E. papayae* JGD233, *E. persicina* NBRC 102418^T^, *E. phyllosphaerae* CMYE1^T^, *E. pyri* DE2^T^, *E. rhapontici* BY21311 and the *Erwinia* sp. PL328 isolated from dogwood **(Supplementary Table 6)**. The levansucrase gene (*lsc*) was only detected in *E. amylovora* ATCC15580^T^ and LMG1877, *E. tasmaniensis* Et1/99^T^ and *E. piriflorinigrans* CFBP5888^T^. Sorbitol metabolism (srl) operon consisting of 7 genes (*srlA*, *srlE*, *srlB*, *srlD*, *srlM*, *srlR* and *srlQ*) was conserved only in *E. amylovora* ATCC15580^T^ and LMG1877, and *E. pyrifoliae* DSM12163^T^ genomes, while the corresponding operon could not be identified in any other *Erwinia* species (**Figure 5E**).

The desferrioxamine siderophore biosynthetic cluster (*dfoA*, *dfoJ* and *dfoC*), one of the iron-chelating systems, was conserved across *E. amylovora* ATCC15580^T^ and LMG1877, *E. pyrifoliae* DSM12163^T^, *E. tasmaniensis* Et1/99^T^ (**Figure 5F**), *E. piriflorinigrans* CFBP5888^T^, and *E. psidii* IBSBF435^T^. Homologs of the *dfo* genes were also identified in *E. aeris* ACC02193^T^, *E. billingae* Eb661, *E. mallotivora* ICMP5705^T^, *E. persicina* ZSR3, *E. oleae* DAPP-PG531^T^*, E. papayae* JGD 233, *E. pyri* DE2^T^, *E. plantamica* OPT-41^T^, *E. tasmaniensis* PPS120, and *E. typographi* MO43b. However, these homologs exhibited variable levels of sequence conservation and did not constitute a complete conserved *dfo* biosynthetic cluster **(Supplementary Table 6)**. The complete *rosA*-*rosG* cluster, responsible for proferrorosamine production and characteristic pink pigmentation, was present only in the dogwood *Erwinia* sp. PL328, *E. rhapontici* BY21311, and *E. persicina* NBRC102418^T^. Although the complete cluster was identified in these three genomes, BLASTp analysis revealed variable amino acid sequence identity and query coverage among the individual ros proteins **(Supplementary Table 6)**. Two distinct sets of genes encoding flagellar biosynthesis and chemotaxis-related proteins were identified in *E. amylovora* ATCC15580^T^ and LMG1877. The first, Flg-1, comprised a complete set of genes distributed across four clusters and was conserved across all *Erwinia* species with varying degrees of similarity, except in *E. endophytica* A41C3 **(Supplementary Figure 3 and Supplementary Table 6)**. The second set flagellar biosynthesis genes, Flg-2, consisted of two clusters, and was found to be highly conserved in *E. amylovora* ATCC15580^T^ and LMG1877, and *E. pyrifoliae* DSM 12163ᵀ. In the remaining *Erwinia* species, BLASTp analysis identified homologs of Flg-2 genes with moderate to low sequence identity relative to *E. amylovora* ATCC15580^T^ **(Supplementary Table 6)**.

## Discussion

Characterized by substantial genomic heterogeneity, broad ecological niches, and unresolved taxonomic relationships [7,84,85], the genus *Erwinia* represents a highly diverse and evolutionary dynamic group within the *Erwiniaceae*.

In this study, comprehensive genomic analyses refined the evolutionary relationship and species boundaries within the genus, identifying previously unrecognized species-level lineages, and revealed extensive variation in gene content and virulence-associated loci. In particular, the dogwood isolate, ‘*Erwinia* sp. PL328’ formed a deeply divergent lineage and was below the accepted species delineation thresholds relative to all currently recognized *Erwinia* species, strongly suggesting that it represents a previously undescribed species-level lineage within the genus [59,86].

The genomic relatedness analyses clearly resolved species-level boundaries within the genus *Erwinia*, with ANI and dDDH analyses identifying several genomically distinct lineages among strains currently assigned to the same species [57,58,59]. Multiple strains formed distinct lineages separate from their expected species clusters, suggesting potential misclassification or previously unrecognized diversity within the genus. These patterns were supported by the core-genome phylogeny, where the phylogenetic tree revealed a well-supported and distinct branching lineage and clades revealing evolutionary separation. Together, these findings underscore the need for continued taxonomic re-evaluation within the genus *Erwinia*, consistent with previous genome-based phylogenomic studies that have highlighted taxonomic complexity and refined species and genus boundaries within *Erwinia* and the family *Erwiniaceae* [86,87]. Although several misclassified strains were identified, we still used their original species names in this study to avoid confusion until formal taxonomic revisions are undertaken.

The pan-genome and ortholog analyses demonstrate that the *Erwinia* genus possesses an open pan-genome, characterized by a relatively small set of conserved core genes, and a large repertoire of cloud genes compared to reports for other Enterobacteriales [36,88,89]. Considering the average genome size across *Erwinia* species (∼3.7–5.5 Mb), the core genome represents only a small proportion of the total gene content, indicating substantial genomic variation across the genus. The high number of cloud genes further suggests that, while a conserved core set of functions is maintained across the genus, each newly incorporated *Erwinia* genome contributes a substantial number of novel gene families, reflecting an extensive accessory genome and high genomic plasticity. This pattern is characteristic of an open pangenome and is consistent with current models of prokaryotic pangenome evolution, in which the continual acquisition of genes through horizontal gene transfer, together with adaptation to diverse ecological niches, promotes gene-content variation and the maintenance of open pangenomes [90,91]. In the genus *Erwinia*, adaptation to diverse ecological niches, including different plant hosts, insect associations, epiphytic lifestyles, and environmental reservoirs, may have contributed to the extensive accessory genome observed in this study. The comparatively higher number of accessory genes observed in most type/representative species of *Erwinia* also aligns with the pan-genome concept described by Tettelin et al. [92], wherein a conserved core genome is complemented by an accessory genome that contributes substantially to genetic diversity among strains. Consistent with this, genome variability is facilitated by horizontal gene transfer and mobile genetic elements, including insertion sequences, plasmids, and phages, which promote changes in gene content and genome plasticity [93]. Similar patterns have been reported in other members of Enterobacterales order, such as *Enterobacter hormaechei* complex, where genome diversification is driven by mobile genetic elements including plasmid replicons, prophages, integrative conjugative elements, and transposable elements [94]. Despite the overall open pan-genome observed at the genus level, not all *Erwinia* species exhibit the same degree of accessory genome diversity. The Rosaceae-associated pathogens *E. amylovora* ATCC15580^T^, *E. pyrifoliae* DSM12163^T^, *E. tasmaniensis* Et1/99^T^, and *E. piriflorinigrans* CFBP5888^T^ possessed comparatively fewer accessory and unique genes, suggesting greater genome conservation associated with host specialization. Consistent with the ortholog analysis, these species also shared the high overlap in orthologous gene content and the few species-specific genes among the lineages examined, indicating conservation of the genomic repertoire associated with bacterial survival, host colonization, and virulence. Adaptation to a relatively restricted host range may have shaped a more conserved gene repertoire through selective gene retention and loss. This observation is consistent with the concept that niche-specialized bacteria tend to possess more closed pan-genomes than species inhabiting diverse ecological niches and more variable environments [82,91]. However, the high number of unique genes in *E. tracheiphila* ICMP5845^T^ and *E. typographi* MO43b likely reflects the evolutionary diversification of lineages occupying different ecological and host-associated environments. In particular, the presence of mobile genetic element- and phage-associated genes in *E. typographi* MO43b suggests increased genome plasticity that may have facilitated adaptation to its insect-associated ecological niche. This interpretation is supported by previous studies of *E. tracheiphila*, which have demonstrated that accessory and horizontally acquired genes contribute directly to host colonization, virulence, and ecological specialization, highlighting the evolutionary significance of accessory genomic regions in shaping niche adaptation within the genus [20,80,95]. In contrast, the non-pathogenic lineages (*E. oleae* DAPP-PG531^T^, *E. tasmaniensis* PPS120, *E. phyllosphaerae* CMYE1^T^, and *E. aeris* ACCC02193^T^) retained relatively few unique genes, indicating a more conserved genomic repertoire and comparatively limited lineage-specific diversification.

Mobile genetic elements further contributed to the genomic diversity observed among *Erwinia* lineages. As mobile elements potentiate gene gain and loss [96], the marked variation in genomic islands and prophage content across species suggests that horizontal gene transfer dynamics are not uniform throughout the genus. Genomic island abundance ranged from fewer than 20 islands in certain species to more than 50 in others, reflecting substantial differences in the acquisition and retention of these horizontally acquired regions that may have contributed to host association, ecological adaptation, and genome diversification within the genus [97]. Likewise, prophage content also varied considerably among species. While some lineages retain multiple prophage remnants, others, including *E. amylovora* ATCC 15580 ^T^, *E. amylovora* LMG 1877, *E. phyllosphaerae* CMYE1^T^, *E. sorbitola* J780^T^, and *E. tasmaniensis* Et1/99^T^ showed an absence of complete and intact phage-derived sequences, reflecting distinct histories of phage invasion, excision, or domestication [98]. Such variation in prophage content may also have contributed to horizontal gene exchange, genome plasticity, ecological adaptation, and the evolution of virulence-associated traits within *Erwinia* [69,99]. The dogwood isolate PL328 contained one intact prophage, several incomplete prophages and no antimicrobial resistance genes. The limited distribution of antimicrobial resistance determinants across the genus indicates that resistance traits have not been widely retained during the evolutionary diversification of the genus and may represent lineage-specific acquisitions associated with localized selective pressures [100].

Virulence in gram-negative bacteria is primarily mediated by type secretion systems, and extracellular polysaccharides that influence the host specificity and pathogenicity [101]. In this study, comparative genomic analyses of key virulence determinants (Type III and Type VI secretion systems, metalloproteases, exopolysaccharides, flagellar cluster, sorbitol metabolism and iron scavenging siderophore) revealed broad variation across species, including pathogenic, non-pathogenic, insect-associated, and environmental species. The major pathogenic species, *E. amylovora* (isolates ATCC15580^T^, LMG1877) and *E. pyrifoliae* DSM12163^T^ shared nearly all core virulence genes, differing only in a few factors such as the inv/spa secretion islands, and the absence of the third T6SS cluster and metalloproteases in *E. pyrifoliae* DSM12163^T^ [32, 72]. Interestingly, these species also exhibited high genomic similarity to the non-pathogenic epiphyte *E. tasmaniensis* Et1/99^T^, supporting previous observations that pathogenic and non-pathogenic pome fruit-associated *Erwinia* species share a conserved genomic background despite their contrasting biology [102]. Despite the conservation of the Hrp-T3SS in both pathogenic and non-pathogenic species, pathogenicity appears to be associated with the presence of additional determinants, including the sorbitol utilization operon, metalloproteases, and the complete amylovoran biosynthetic cluster, rather than the Hrp-T3SS alone [38,73,103]. Interestingly, the amylovoran biosynthetic locus also exhibited a lineage-specific distribution. Complete *ams/cps* clusters were retained in the *E. amylovora, E. pyrifoliae,* and *E. tasmaniensis* lineages, whereas no *ams* homologs were detected in the distinct *E. persicina* ZSR3, *E. aeris* ACC02193^T^, and *E. plantamica* OPT-41^T^ lineage. The concordance between the distribution of the amylovoran biosynthetic cluster and the phylogenomic relationships identified in this study suggests that this major virulence determinant has been evolutionary conserved within specific lineages rather than being uniformly distributed across the genus.

Other plant pathogenic species including *E. piriflorinigrans* CFBP5888^T^, *E. mallotivora* ICMP5705^T^, *E. psidii* IBSBF435^T^, *E. tracheiphila* ICMP5845^T^ and *E. papayae* JGD 233 retained substantial similarity to the hrp T3SS like *E. amylovora* ATCC15580^T^ and LMG1877. In contrast, several phytopathogenic species such as *E. persicina* (NBRC 102418^T^, CFBP8797, CFBP8803, ZSR3), *E. pyri* DE2^T^, and *E. rhapontici* BY21311, *E. billingae* Eb661, cause disease despite lacking the conserved pathogenicity determinants typical of fire-blight pathogens (5,13,14,17, 39, 95,104,105,106]. These species appear to have evolved alternative pathogenicity strategies or rely on host-specific ecological contexts that do not require the classical virulence factors (PAI-1, PAI-2 and PAI-3). Together, these patterns indicate that pathogenicity in *Erwinia* is not based on a single conserved set of virulence proteins. Instead, different species secrete distinct combinations of proteins, reflecting diverse evolutionary pathways and host interactions [107]. Consistent with this pattern of lineage-specific accessory determinants, the proferrorosamine biosynthesis (*rosA*-*rosG*) cluster was restricted to a subset of closely related *Erwinia* lineages, being detected only in *E. persicina* NBRC102418^T^, *E. rhapontici* BY21311, and the dogwood *Erwinia* sp. PL328. The *ros* cluster, originally characterized in *E. rhapontici* P45, encodes the siderophore proferrorosamine and has previously been reported in *E. rhapontici* and *E. persicina* [5,108]. Its detection in the *Erwinia* sp. PL328 isolated from dogwood reported in this study expands the known distribution of this locus within the genus and suggests that it has been retained in selected evolutionary lineages rather than being widely conserved across *Erwinia*. These findings highlight the need for further investigation to elucidate the contribution of individual secreted proteins to pathogenicity and host specificity for different species of *Erwinia* genus.

The diversity of virulence-associated gene repertoires extended beyond phytopathogenic species. The insect-associated species, *E. typographi* MO43b, isolated from bark beetles, is non-phytopathogenic [27], whereas *E. aphidicola* JCM21238^T^ and *E. sorbitola* J780^T^, initially recovered from insects or animal sources retain sufficient virulence-associated determinants to infect plants [26,109]. However, their virulence profiles differ substantially from the canonical fire blight-associated determinants of *E. amylovora*. Notably, although Tao et al. [26] reported a complete T6SS in *E. sorbitola* J780^T^, the corresponding locus could not be recovered using the *E. amylovora* T6SS proteins as BLASTp queries in the present study, suggesting that the T6SS of *E. sorbitola* is highly divergent from that of *E. amylovora*. Together, these observations indicate that insect-associated *Erwinia* species possess distinct virulence repertoires that differ from the canonical pathogenicity determinants of fire blight pathogens. Environmental and endophytic species such as *E. oleae* DAPP-PG531^T^ from olive knots [23], *E. phyllosphaerae* CMYE1^T^ from the pomelo phyllosphere [24], *E. aeris* ACC02193^T^ from surface of ore [28], *E. endophytica* A41C3 from potato [22] and *E. plantamica* OPT-41^T^ [110] possessed only weak remnants of virulence-associated loci or lacked them entirely. This observation further supports previous comparative genomic studies showing that closely related pathogenic and non-pathogenic *Erwinia* species differ primarily in accessory virulence determinants while sharing a conserved genomic background [102]. Overall, genomic analyses indicate that the distribution of virulence-associated determinants across the genus *Erwinia* reflects both phylogenetic relationships and ecological specialization. Closely related species generally share a conserved genomic backbone, whereas differences in accessory virulence determinants are associated with adaptation to distinct ecological niches, consistent with patterns reported in other members of the family *Erwiniaceae*. Similarly, comparative genomic analyses of the closely related genus *Pantoea* have shown that adaptation to phytopathogenic, endophytic, environmental, and clinical lifestyles is accompanied by lineage-specific differences in accessory genes involved in host interaction, virulence, and niche adaptation [111]. Likewise, *Erwinia* species closely associated with plant hosts, including *E. amylovora*, *E. pyrifoliae*, and *E. piriflorinigrans*, retained a larger complement of canonical virulence determinants, whereas insect-associated, environmental, and endophytic species generally possessed only subsets of these loci or lacked several of them entirely. Together, these findings suggest that diversification within *Erwinia* has been shaped by lineage-specific variation in accessory virulence determinants, reflecting evolutionary adaptation to distinct ecological niches while maintaining a conserved genomic backbone.

When considered together with the ANI/dDDH analyses, core-genome phylogeny, pan-genome structure, and lineage-specific distribution of mobile genetic elements and virulence determinants, our findings demonstrate that *Erwinia* is a genomically diverse genus undergoing continual evolutionary diversification. Rather than relying on a universally conserved virulence repertoire, pathogenicity appears to have evolved through lineage-specific combinations of accessory genes shaped by evolutionary history and ecological adaptation. These findings provide a comprehensive genomic framework for refining *Erwinia* taxonomy, improving the prediction of pathogenic factors, and advancing our understanding of the evolutionary processes underlying diversification within the genus.

## Funding information

This research was supported by the United States Department of Agriculture, National Institute of Food and Agriculture, Award No. 2023-67013-39301and also by Dr. Mohammad Arif’s Hatch Project at Oklahoma State University.

The mention of trade names or commercial products in this publication is solely for the purpose of providing specific information and does not imply recommendation or endorsement by Oklahoma State University.

## Supplementary Figures and Tables

**Supplementary Figure 1.** Pan- and core-genome accumulation curves of 104 *Erwinia* genomes. The dashed blue line shows the expansion of the total number of genes (pan-genome) as each of the 104 genomes is added, reflecting the genomic diversity within the genus. The solid red line represents the number of genes conserved across all genomes (core genome). The pan-core genome analysis was performed using Roary v3.13.0, and the curve were plotted in R. These results indicate that *Erwinia* possesses an open pan-genome, with new genomes continually contributing novel gene content.

**Supplementary Figure 2.** Orthologous gene cluster analysis of the main *Erwinia* clade, including the sister clade *Erwinia aphidicola* JCM 21238^T^.

**Supplementary Figure 3.** Distribution and comparative organization of flagellar gene clusters among *Erwinia* species identified by reference mapping in Geneious Prime v 2025.2. **(A)** Flagellar cluster I (Flg1), **(B)** Flagellar cluster II (Flg2)

**Supplementary Table 1.** List of *Erwinia* strains included in the core-genome phylogenetic analysis, including species designation, host of isolation, geographic origin, and genome accession information.

**Supplementary Table 2**. Selected *Erwinia* genomes used for comparative genomic analyses. Type strains were included whenever available; otherwise, representative strains were selected for species lacking an available type strain genome.

**Supplementary Table 3**. Genomic islands predicted in the type and representative genomes of Erwinia species using IslandViewer4. Each worksheet corresponds to a single genome and lists the predicted genomic islands, including their genomic coordinates, size, prediction method(s), and annotated gene content.

**Supplementary Table 4.** Prophage regions identified in *Erwinia* strains using PHASTER. The table includes the genome, prophage region number, region length, completeness classification, PHASTER score, predicted phage keyword, genomic position, total number of predicted proteins, and the numbers of phage-hit, hypothetical, and bacterial proteins within each prophage region.

**Supplementary Table 5.** Antimicrobial resistance genes identified in *Erwinia persicina* CFBP 8797 and NBRC 102418^T^.

**Supplementary Table 6.** Percent identity and query coverage of T3SS-, T6SS-, and other virulence-associated proteins identified by BLASTP using *Erwinia amylovora* ATCC15580T as the reference against all type and representative *Erwinia* species.

