## Supplementary for "Phylogenomics and comparative genomics of the genus *Erwinia* reveal taxonomic inconsistencies and evolutionary diversification": Supplementary Table 1.docx

**Supplementary Table 1.** List of *Erwinia* strains included in the core-genome phylogenetic analysis, including species designation, host of isolation, geographic origin, and genome accession information.

| **Sl. No.** | **Scientific name** | **Strain** | **GenBank** | **Genome size (Mb)** | **Level** | **GC %** | **% Completeness** | **% Contamination** | **Collection Year** | **Geographic Location** | **Isolation source** |
| --- | --- | --- | --- | --- | --- | --- | --- | --- | --- | --- | --- |
| 1 | *Erwinia aeris* | ACCC 02193^T^ | GCA_041224955.1 | 4.96 | Contig | 55.5 | 95.61 | 5.02 | 2005 | China | Ore |
| 2 | *Erwinia amylovora* | EaSmR | GCA_043228865.1 | 3.83 | Complete | 53.5 | 98.04 | 0.22 | 2022 | South Korea | Cell culture |
| 3 | *Erwinia amylovora* | ATCC 15580^T^ | GCA_017161565.1 | 3.84 | Complete | 53.5 | 95.4 | 0.2 | 2020 | United Kingdom | *Pyrus communis* |
| 4 | *Erwinia amylovora* | Ea1/79Sm | GCA_015650045.1 | 3.84 | Complete | 53.5 | 98.23 | 0.2 | 1979 | Germany | *Malus sylvestris* |
| 5 | *Erwinia amylovora* | Ea 1/79 | GCA_040125995.1 | 3.84 | Complete | 53.5 | 98.23 | 0.2 | 1979 | Germany | *Malus domestica* |
| 6 | *Erwinia amylovora* | E-2 | GCA_002803865.1 | 3.84 | Complete | 53.5 | 98.23 | 0.2 | 2007 | Belarus | *Malus* sp. |
| 7 | *Erwinia amylovora* | CP200242 | GCA_027557715.1 | 3.83 | Complete | 53.5 | 98.21 | 0.2 | 2020 | South Korea | Asian pear |
| 8 | *Erwinia amylovora* | FB-86 | GCA_012980785.1 | 3.83 | Complete | 53.5 | 98.21 | 0.2 | 2015 | South Korea | Apple tree |
| 9 | *Erwinia amylovora* | PBI209 | GCA_040105085.1 | 3.83 | Complete | 53.5 | 96.43 | 0.76 | 2023 | China | Pear |
| 10 | *Erwinia amylovora* | Ea1189 | GCA_016446415.2 | 3.83 | Complete | 53.5 | 97.03 | 0.21 | 2018 | Germany | Apple |
| 11 | *Erwinia amylovora* | ATCC 49946 | GCA_000027205.1 | 3.91 | Complete | 53.5 | 98.23 | 0.2 | - | United Kingdom | Malus |
| 12 | *Erwinia amylovora* | CFPB1430 | GCA_000091565.1 | 3.83 | Complete | 53.5 | 97.64 | 0.22 | 1972 | France | *Crataegus* |
| 13 | *Erwinia amylovora* | FC01 | GCA_042055245.1 | 3.96 | Contig | 53.5 | 96.67 | 0.56 | 2023 | USA | Apple orchards |
| 14 | *Erwinia amylovora* | Ea321 | GCA_003992295.1 | 3.87 | Contig | 53.5 | 97.97 | 1.87 | 1972 | France:Nord | Hawthorn |
| 15 | *Erwinia amylovora* | Fb-97b | GCA_012371505.1 | 3.80 | Contig | 53.5 | 96.86 | 0.2 | 1993 | New Zealand | *Malus domestica* |
| 16 | *Erwinia amylovora* | Ea650 | GCA_012367155.1 | 3.77 | Contig | 53.5 | 98.36 | 0.24 | 1993 | Poland | *Crataegus monogyna* |
| 17 | *Erwinia amylovora* | E22 | GCA_040234495.1 | 3.80 | Complete | 53.5 | 95.97 | 0.76 | 2022 | Kazakhstan | *Malus* spp. |
| 18 | *Erwinia amylovora* | 7-3 | GCA_020544325.1 | 3.99 | Chromosome | 53.5 | 97 | 1.77 | 2019 | USA:Califonia | *Malus domestica* |
| 19 | *Erwinia amylovora* | HKN06P1 | GCA_004023365.1 | 3.90 | Contig | 53.5 | 98.21 | 1.62 | 2006 | USA:Pennsylvania | Apple tree |
| 20 | *Erwinia amylovora* | Ea7-96 | GCA_012371615.1 | 3.81 | Contig | 53.5 | 93.13 | 0.92 | 1996 | Canada | *Rubus idaeus* |
| 21 | *Erwinia amylovora* | CEB_13 | GCA_050751315.1 | 3.84 | Contig | 53.5 | 98.38 | 0.26 | 2024 | Portugal | Soil |
| 22 | *Erwinia amylovora* | 57743_ID1772_81-CP_06_S121_L001 | GCA_023182895.1 | 3.81 | Scaffold | 53.5 | 97.66 | 0.27 | 2020 | Italy | *Cydonia oblonga* |
| **23** | ***Erwinia amylovora*** | **LMG1877** |  | **3.83** | **Complete** | **53.5** |  |  | **1972** | **Denmark** | ***Cydonia oblonga*** |
| 24 | *Erwinia aphidicola* | B09_9 | GCA_036866135.1 | 4.701 | Complete | 56.5 | 92.94 | 3.06 | 2021 | Denmark | Flag leaf of wheat |
| 25 | *Erwinia aphidicola* | USMM130 | GCA_037149315.1 | 4.838 | Contig | 56.5 | 98.45 | 4.03 | 2012 | USA: Pennsylvania | Mosquito |
| 26 | *Erwinia aphidicola* | 1B06 | GCA_050038595.1 | 4.88 | Complete | 57 | 98.45 | 5.35 | 2012 | USA: California | *Lactuca sativa* |
| 27 | *Erwinia aphidicola* | W09_2 | GCA_036865745.1 | 4.653 | Complete | 56.5 | 93.4 | 4.27 | 2021 | Denmark | Flag leaf of wheat |
| 28 | *Erwinia aphidicola* | QCJ3A | GCA_037045795.1 | 5.16 | Contig | 56.5 | 98.51 | 5.82 | 2021 | Chile | *Allium cepa* |
| 29 | *Erwinia aphidicola* | h-5 TE3250 | GCA_050435985.1 | 4.796 | Contig | 57 | 97.55 | 3.87 | - | - | - |
| 30 | *Erwinia aphidicola* | QCJ2B | GCA_037045765.1 | 5.257 | Contig | 56.5 | 98.8 | 5.81 | 2021 | Chile | *Allium cepa* |
| 31 | *Erwinia aphidicola* | EPB1 | GCA_037045825.1 | 5.407 | Contig | 56 | 98.79 | 6.27 | 2023 | Chile | *Allium cepa* |
| 32 | *Erwinia aphidicola* | JCM 21238^T^ | GCA_014773485.1 | 5.118 | Contig | 56.5 | 98.77 | 12.93 | 1996 | Japan | Pea aphid |
| 33 | *Erwinia aphidicola* | USHLN281 | GCA_037149555.1 | 4.837 | Contig | 56.5 | 98.45 | 4.03 | 2012 | USA: Pennsylvania | Mosquito |
| 34 | *Erwinia aphidicola* | Bi13 | GCA_918698235.1 | 4.939 | Contig | 56.5 | 98.45 | 5.1 | 2017 | United Kingdom | Barely rhizosphere |
| 35 | *Erwinia aphidicola* | USHLN33 | GCA_037144385.1 | 4.905 | Contig | 56.5 | 98.74 | 4.9 | 2016 | USA: Florida | Mosquito |
| 36 | *Erwinia aphidicola* | Z9_3 | GCA_036865565.1 | 4.641 | Complete | 56.5 | 93.45 | 3.26 | 2021 | Denmark | Flag leaf of wheat |
| 37 | *Erwinia billingiae* | Eb661 | GCA_000196615.1 | 5.372 | Complete | 55 | 99.73 | 0.19 | 1959 | England | *Pyrus* spp. |
| 38 | *Erwinia billingiae* | TH88 | GCA_008728215.1 | 4.903 | Complete | 55 | 98.78 | 0.7 | 2017 | USA: Alaska | Soil core |
| 39 | *Erwinia billingiae* | W05_1 | GCA_036865555.1 | 4.791 | Complete | 55 | 95.8 | 0.7 | 2021 | Denmark | Flag leaf of wheat |
| 40 | *Erwinia billingiae* | Eb21-1 | GCA_039675705.1 | 5.136 | Scaffold | 55 | 99.13 | 0.52 | 1997 | Canada | *Pyrus communis* |
| 41 | *Erwinia billingiae* | MYb121 | GCA_037479335.1 | 5.027 | Scaffold | 55 | 97.97 | 1.09 | 2020 | Germany | *Caenorhabditis elegans* |
| 42 | *Erwinia billingiae* | LS-1 | GCA_026233155.1 | 5.123 | Scaffold | 55 | 97.15 | 0.71 | 2018 | China | *Gastrodia elata* |
| 43 | *Erwinia billingiae* | PBb | GCA_017168115.1 | 5.259 | Scaffold | 55 | 98.55 | 1.31 | 2010 | New Zealand | *Pyrus* spp. |
| 44 | *Erwinia billingiae* | OSU19-1 | GCA_001269445.1 | 5.602 | Scaffold | 55 | 91.08 | 11.96 | 2015 | USA: Oregon | *Pyrus communis* |
| 45 | *Erwinia endophytica* | A41C3 | GCA_009295515.1 | 4.227 | Scaffold | 51.5 | 85.99 | 3.48 | 2009 | Portugal | *Pinus pinaster* |
| 46 | *Erwinia mallotivora* | BT-MARDI | GCA_000590885.1 | 4.64 | Contig | 52.5 | 88.02 | 4.99 | 2010 | Malaysia | Papaya tree |
| 47 | *Erwinia mallotivora* | ICMP 5705^T^ | GCA_042432085.1 | 4.409 | Contig | 52.5 | 88.92 | 2.97 | 2020 | Japan | *Mallotus japonicus* |
| 48 | *Erwinia mallotivora* | EP21 | GCA_025388925.1 | 4.896 | Scaffold | 52.5 | 93.39 | 5.73 | 2014 | Philippines | Papaya petiole |
| 49 | *Erwinia mallotivora* | EP61 | GCA_025388785.1 | 4.967 | Scaffold | 52.5 | 93.6 | 5.8 | 2012 | Philippines | Papaya stem |
| 50 | *Erwinia mallotivora* | EP60 | GCA_025388825.1 | 4.994 | Scaffold | 52.5 | 93.77 | 5.89 | 2014 | Philippines | Papaya stem |
| 51 | *Erwinia oleae* | DAPP-PG531^T^ | GCA_000770305.1 | 4.744 | Contig | 54.5 | 92.24 | 2.96 | 2003 | Italy | Olive knot |
| 52 | *Erwinia papayae* | JGD 233 | GCA_040741485.1 | 4.957 | Contig | 52.5 | 92.96 | 5.12 | 2023 | Guam | *Carica papaya* |
| 53 | *Erwinia persicina* | ZSR3 | GCA_024168495.1 | 4.695 | Contig | 56 | 95.68 | 4.61 | - | USA: Monroe | - |
| 54 | *Erwinia persicina* | DOAB1061 | GCA_014357465.1 | 4.892 | Contig | 55.5 | 96.22 | 7.73 | 2017 | Canada | Wheat |
| 55 | *Erwinia persicina* | CFBP8797 | GCA_014838825.1 | 4.806 | Contig | 56 | 96.69 | 5.53 | 2016 | France | *Raphanus sativus* seed |
| 56 | *Erwinia persicina* | CFBP8803 | GCA_014839105.1 | 4.897 | Contig | 56 | 96.69 | 5.37 | 2016 | France | *Raphanus sativus* flower |
| 57 | *Erwinia persicina* | SR13 | GCA_024436315.1 | 4.806 | Scaffold | 55.5 | 96.54 | 7.16 | 2019 | USA: Iowa | Asparagus |
| 58 | *Erwinia persicina* | CFBP8767 | GCA_014839215.1 | 5.182 | Contig | 55 | 97.47 | 6.43 | 2017 | France | *Brassica napus* seed |
| 59 | *Erwinia persicina* | CFBP13511 | GCA_005233475.1 | 5.203 | Contig | 55 | 97.38 | 7.02 | 2014 | France | *Raphanus sativus* seed |
| 60 | *Erwinia persicina* | SR16 | GCA_024436235.1 | 4.869 | Scaffold | 55.5 | 96.9 | 6.46 | 2019 | USA: Iowa | Green onion |
| 61 | *Erwinia persicina* | CFBP13732 | GCA_014839305.1 | 5.183 | Contig | 55 | 97.47 | 6.43 | 2017 | France | *Brassica napus* seed |
| 62 | *Erwinia persicina* | Cp2 | GCA_019844095.1 | 4.803 | Complete | 55.5 | 96.4 | 6.1 | 2019 | China | Alfalfa seeds |
| 63 | *Erwinia persicina* | B64 | GCA_003485445.1 | 5.07 | Complete | 55 | 95.93 | 6.84 | 2016 | South Korea | *Allium cepa* |
| 64 | *Erwinia persicina* | SR15 | GCA_024397315.1 | 4.898 | Complete | 55.5 | 96.72 | 6.22 | 2019 | USA: Iowa | Green onion |
| 65 | *Erwinia persicina* | ML2-2023-5 | GCA_037081855.1 | 5.014 | Complete | 55.5 | 96.76 | 6.77 | 2023 | Australia | Leaf wash from mixed-leaf salad |
| 66 | *Erwinia persicina* | BST187 | GCA_047302665.1 | 4.852 | Complete | 55.5 | 96.4 | 6.36 | 2022 | China | Rhizosphere soil |
| 67 | *Erwinia persicina* | NBRC 102418^T^ | GCA_001571305.1 | 4.906 | Contig | 55.5 | 96.69 | 5.68 | Unknown | Japan | Tomato |
| 68 | *Erwinia phyllosphaerae* | CMYE1^T^ | GCA_019132875.1 | 4.732 | Contig | 54 | 95.62 | 4.59 | 2021 | China | Citrus phyllosphere |
| 69 | *Erwinia piriflorinigrans* | CFBP 5888^T^ | GCA_001050515.1 | 3.931 | Contig | 53 | 97.68 | 0.78 | 1999 | Spain | Pear blossoms |
| 70 | *Erwinia plantamica* | OPT-41^T^ | GCA_043420595.1 | 4.764 | Contig | 55.5 | 95.69 | 5.16 | 2019 | Russia | Seedlings of spring wheat |
| 71 | *Erwinia psidii* | IBSBF 435^T^ | GCA_003846135.1 | 4.503 | Scaffold | 51.5 | 89.27 | 4.14 | 1987 | Brazil | *Psidium guajava* |
| 72 | *Erwinia psidii* | LPF 534 | GCA_026549045.1 | 4.477 | Scaffold | 51.5 | 89.34 | 4.17 | 2016 | Brazil | *Eucalyptus dunnii* |
| 73 | *Erwinia psidii* | LPF 681 | GCA_026549085.1 | 4.667 | Scaffold | 51.5 | 89.21 | 3.92 | 2016 | Brazil | *Psidium guajava* |
| 74 | *Erwinia psidii* | LPF 640 | GCA_026549055.1 | 4.542 | Scaffold | 51.5 | 89.15 | 4.1 | 2016 | Brazil | *Eucalyptus* sp. |
| 75 | *Erwinia pyri* | DE2^T^ | GCA_030758455.1 | 4.758 | Complete | 54.5 | 94.76 | 4.17 | 2022 | China | *Pyrus* orchard |
| 76 | *Erwinia pyrifoliae* | EpK1/15 | GCA_002952315.1 | 4.076 | Complete | 53.5 | 95.79 | 0.59 | 2017 | South Korea | Apple twig |
| 77 | *Erwinia pyrifoliae* | DSM 12163^T^ | GCA_000026985.1 | 4.073 | Chromosme | 53.5 | 97.08 | 0.41 | 1996 | South Korea | *Pyrus* spp. |
| 78 | *Erwinia pyrifoliae* | CP201486 | GCA_025159015.1 | 4.142 | Complete | 53.5 | 95.59 | 2.46 | 2020 | South Korea | Apple Orchard |
| 79 | *Erwinia pyrifoliae* | YKB12327 | GCA_041015235.1 | 4.062 | Complete | 53.5 | 84.68 | 1.1 | 2015 | South Korea | *Malus domestica* |
| 80 | *Erwinia pyrifoliae* | Ep1/96 | GCA_000027265.1 | 4.073 | Complete | 53.5 | 97.35 | 0.41 | 1996 | South Korea | *Pyrus* spp. |
| 81 | *Erwinia pyrifoliae* | YKB12328 | GCA_020428355.1 | 4.079 | Contig | 53.5 | 96.72 | 0.41 | 2015 | South Korea | Apple twig |
| 82 | *Erwinia pyrifoliae* | CP20113301 | GCA_025402895.1 | 4.161 | Complete | 53.5 | 95.59 | 2.53 | 2020 | South Korea | Apple |
| 83 | *Erwinia rhapontici* | BY21311 | GCA_020683125.1 | 5.165 | Complete | 54 | 98.01 | 6.12 | 2021 | China | *Apium graveolens* |
| 84 | *Erwinia rhapontici* | 1SR | GCA_050613375.1 | 5.439 | Complete | 54 | 95.12 | 6.89 | 2018 | Chile | Soil |
| 85 | *Erwinia rhapontici* | MAFF 311154 | GCA_018326325.1 | 5.36 | Complete | 54 | 93.57 | 4.03 | 1985 | Japan | *Brassica rapa* |
| 86 | *Erwinia rhapontici* | WS3235 | GCA_017875455.1 | 5.454 | Contig | 54 | 98.01 | 6.82 | - | - | - |
| 87 | *Erwinia rhapontici* | BIGb0389 | GCA_024807835.1 | 5.399 | Contig | 54 | 98.01 | 6.23 | - | - | - |
| 88 | *Erwinia rhapontici* | CGMCC1.6978 | GCA_009846845.1 | 5.488 | Contig | 54 | 98.01 | 8.52 | 1924 | Germany | Plant |
| 89 | *Erwinia rhapontici* | QL1 | GCA_049206705.1 | 5.153 | Scaffold | 54 | 98.01 | 6.3 | 2019 | Canada | *Vaccinium corymbosum* |
| 90 | *Erwinia rhapontici* | H1 | GCA_012271765.1 | 5.24 | Scaffold | 54 | 97.79 | 7.07 | - | Hungary | *Prunis persica* |
| 91 | *Erwinia rhapontici* | BIGb0435 | GCA_004364855.1 | 5.288 | Scaffold | 54 | 97.83 | 5.7 | - | - | - |
| 92 | *Erwinia rhapontici* | EDr1-9 | GCA_049199025.1 | 5.102 | Scaffold | 54.5 | 97.08 | 6.27 | 2019 | Canada | *Vaccinium corymbosum* |
| 93 | *Erwinia rhapontici* | MAFF 311153 | GCA_018409035.1 | 5.233 | Complete | 54 | 97.58 | 5.41 | 1985 | Japan | *Brassica rapa* |
| 94 | *Erwinia sorbitola* | J780^T^ | GCA_009738185.1 | 4.75 | Complete | 53 | 97.56 | 5.18 | 2018 | China | Ruddy shelduck |
| 95 | *Erwinia tasmaniensis* | Et1/99^T^ | GCA_000026185.1 | 4.068 | Complete | 53.5 | 99.62 | 0.03 | 1999 | Tasmania, Australia | Apple flower |
| 96 | *Erwinia tasmaniensis* | PPS120 | GCA_047918655.1 | 4.561 | Scaffold | 54.5 | 95.56 | 1.92 | 2022 | USA: Alabama | Environmental sample |
| 97 | *Erwinia tracheiphila* | BHKY | GCA_021365465.1 | 4.959 | Complete | 50.5 | 79.67 | 2.24 | 2010 | USA:Kentucky | *Cucurbita moschata* |
| 98 | *Erwinia tracheiphila* | BuffGH | GCA_021365505.1 | 4.979 | Complete | 50.5 | 78.89 | 2.24 | 2009 | USA: Pennsylvania | *Cucurbita pepo* ssp. *texana* |
| 99 | *Erwinia tracheiphila* | MDCuke | GCA_021365535.1 | 5.054 | Complete | 50.5 | 79.67 | 2.27 | 2010 | USA: Maryland | *Cucumis sativus* |
| 100 | *Erwinia tracheiphila* | SCR3 | GCA_021365485.1 | 4.975 | Complete | 50.5 | 79.12 | 2.24 | 2009 | USA: Iowa | *Cucumis melo* |
| 101 | *Erwinia tracheiphila* | PSU-1 | GCA_000404125.1 | 4.717 | Scaffold | 50 | 78.86 | 2.15 | - | USA | Infected cucurbits |
| 102 | *Erwinia tracheiphila* | ICMP 5845^T^ | GCA_042427805.1 | 4.935 | Contig | 50.5 | 47.09 | 0.32 | 2022 | USA | *Cucumis* |
| 103 | *Erwinia typographi* | M043b | GCA_000773975.1 | 5.75 | Contig | 55 | 95.65 | 4.98 | 2013 | Malaysia | Waterfall |
| **104** | ***Erwinia* spp.** | **PL328** |  | **5.12** | **Complete** | **53.5** |  |  | **1989** | **USA:Maryland** | ***Cornus florida*** |
