## Supplementary for "Phylogenomics and comparative genomics of the genus *Erwinia* reveal taxonomic inconsistencies and evolutionary diversification": Supplementary Table 2.docx

**Supplementary Table 2.** Selected *Erwinia* genomes used for comparative genomic analyses. Type strains were included whenever available; otherwise, representative strains were selected for species lacking an available type strain genome.

| **Sl. No.** | **Pathogen** | **Genbank accession number** | **Genome completeness** | **Feature** |
| --- | --- | --- | --- | --- |
| 1 | *Erwinia aeris* ACC02193^T^ | GCA_041224955.1 | Contig | Environment isolated |
| 2 | *Erwinia amylovora* ATCC15580^T^ | GCA_017161565.1 | Complete | Plant pathogenic |
| 3 | *Erwinia amylovora* LMG1877 |  |  | Plant pathogenic** |
| 4 | *Erwinia aphidicola* JCM21238^T^ | GCA_014773485.1 | Contig | Plant pathogenic and insect associated |
| 5 | *Erwinia billingiae* Eb661 | GCA_000196615.1 | Complete | Plant and human associated |
| 6 | *Erwinia endophytica* A41C3 | GCA_009295515.1 | Scaffold | Plant associated |
| 7 | *Erwinia mallotivora* ICMP5705^T^ | GCA_042432085.1 | Contig | Plant pathogenic |
| 8 | *Erwinia oleae* DAPP-PG531^T^ | GCA_000770305.1 | Contig | Plant associated |
| 9 | *Erwinia papayae* JGD 233 | GCA_040741485.1 | Contig | Plant pathogenic |
| 10 | *Erwinia piriflorinigrans* CFBP5888^T^ | GCA_001050515.1 | Contig | Plant pathogenic |
| 11 | *Erwinia psidii* IBSBF435^T^ | GCA_003846135.1 | Scaffold | Plant pathogenic |
| 12 | *Erwinia persicina* NBRC 102418^T^ | GCA_001571305.1 | Contig | Plant pathogenic |
| 13 | *Erwinia persicina* CFBP8797 | GCA_014838825.1 | Contig | Plant pathogenic |
| 14 | *Erwinia persicina* CFBP8803 | GCA_014839105.1 | Contig | Plant pathogenic |
| 15 | *Erwinia persicina* ZSR3 | GCA_024168495.1 | Contig | Plant pathogenic |
| 16 | *Erwinia plantamica* OPT-41^T^ | GCA_043420595.1 | Contig | Plant associated* |
| 17 | *Erwinia pyri* DE2^T^ | GCA_030758455.1 | Complete | Plant pathogenic* |
| 18 | *Erwinia pyrifoliae* DSM12163^T^ | GCA_000026985.1 | Complete | Plant pathogenic |
| 19 | *Erwinia phyllosphaerae* CMYE1^T^ | GCA_019132875.1 | Contig | Plant associated |
| 20 | *Erwinia rhapontici* BY21311 | GCA_020683125.1 | Complete | Plant pathogenic |
| 21 | *Erwinia sorbitola* J780^T^ | GCA_009738185.1 | Complete | Plant pathogenic* |
| 22 | *Erwinia* sp. PL328 |  |  | Plant pathogenic** |
| 23 | *Erwinia tracheiphila* ICMP5845^T^ | GCA_042427805.1 | Contig | Plant pathogenic |
| 24 | *Erwinia tasmaniensis* Et1/99^T^ | GCA_000026185.1 | Complete | Plant associated |
| 25 | *Erwinia tasmaniensis* PPS120 | GCA_047918655.1 | Scaffold | Plant associated |
| 26 | *Erwinia typographi* MO43b | GCA_000773975.1 | Contig | Plant and insect associated |

*: Not validly published

**: Sequenced in lab
