## Supplementary for "Phylogenomics and comparative genomics of the genus *Erwinia* reveal taxonomic inconsistencies and evolutionary diversification": Supplementary Table 4.docx

**Supplementary Table 4.** Prophage regions identified in *Erwinia* strains using PHASTER. The table includes the genome, prophage region number, region length, completeness classification, PHASTER score, predicted phage keyword, genomic position, total number of predicted proteins, and the numbers of phage-hit, hypothetical, and bacterial proteins within each prophage region.

| **Genome** | **Region** | **Length** | **Completeness** | **Score** | **Specific Keyword** | **Position** | **Total protein** | **Phage Hit Proteins** | **Hypothetical Proteins** | **Bacterial Proteins** |
| --- | --- | --- | --- | --- | --- | --- | --- | --- | --- | --- |
| *Erwinia amylovora* LMG1877 | 1 | 8.5 Kb | incomplete | 50 | integrase, tail | 29557-38127 | 13 | 8 | 5 | 0 |
|  | 2 | 27.6 Kb | incomplete | 40 | plate, tail, integrase | 2304680-2332289 | 22 | 14 | 8 | 0 |
|  | 3 | 23.2 Kb | incomplete | 50 | tail, integrase | 2366180-2389437 | 8 | 6 | 2 | 0 |
| *Erwinia amylovora* ATCC15580^T^ | 1 | 8.5 Kb | incomplete | 30 | tail, integrase | 1897784-1906300 | 12 | 9 | 3 | 0 |
|  | 2 | 23.2 Kb | incomplete | 40 | tail, recombinase | 3353895-3377152 | 7 | 5 | 0 | 2 |
|  | 3 | 29.6 Kb | incomplete | 60 | envelope, integrase, flippase, tail, plate | 3405970-3435593 | 18 | 12 | 3 | 3 |
| *Erwinia aeris* ACC02193^T^ | 1 | 50.8 Kb | incomplete | 60 | integrase, tail | 164573-215460 | 76 | 54 | 22 | 0 |
|  | 2 | 35.6 Kb | intact | 150 | integrase, portal, terminase, capsid, head, lysis, tail, plate | 1584127-1619746 | 51 | 42 | 9 | 0 |
|  | 3 | 16 Kb | incomplete | 60 | tail, plate, lysis | 2866378-2882444 | 22 | 19 | 3 | 0 |
|  | 4 | 41 Kb | incomplete | 50 | tail, integrase, head | 2886473-2927479 | 26 | 18 | 8 | 0 |
|  | 5 | 11.6Kb | incomplete | 20 | integrase | 3764253-3775897 | 17 | 7 | 10 | 0 |
|  | 6 | 11.9 Kb | incomplete | 40 | integrase, head | 4097654-4109637 | 11 | 8 | 3 | 0 |
|  | 7 | 13.9 Kb | incomplete | 10 | NA | 4250762-4264690 | 24 | 15 | 9 | 0 |
|  | 8 | 39.8 Kb | incomplete | 40 | tail | 4562501-4602392 | 41 | 26 | 15 | 0 |
| *Erwinia aphidicola* JCM21238^T^ | 1 | 13.5 Kb | intact | 130 | tail, plate, lysis, lysin | 1295487-1309011 | 21 | 21 | 0 | 0 |
|  | 2 | 51.6 Kb | intact | 150 | flippase, tail, head, capsid, portal, terminase, lysin, integrase | 3552470-3604149 | 60 | 50 | 10 | 0 |
|  | 3 | 11.6 Kb | incomplete | 30 | integrase, head | 4066026-4077656 | 12 | 7 | 5 | 0 |
| *Erwinia billingae* Eb661 | 1 | 37.4 Kb | intact | 150 | integrase, portal, terminase, capsid, head, lysin, tail, plate | 102203-139664 | 50 | 37 | 13 | 0 |
|  | 2 | 35.6 Kb | questionable | 70 | protease, plate, tail | 1245185-1280792 | 48 | 40 | 8 | 0 |
|  | 3 | 23.8 Kb | questionable | 70 | integrase, tail, plate, protease | 3521948-3545777 | 15 | 10 | 5 | 0 |
|  | 4 | 33.7 Kb | intact | 130 | lysin, portal, terminase, head, capsid, tail, transposase | 3552551-3586339 | 60 | 45 | 15 | 21 |
|  | 5 | 15.2 Kb | incomplete | 20 | tail | 3801028-3816254 | 18 | 12 | 6 | 0 |
|  | 6 | 16.1 Kb | incomplete | 20 | integrase | 4402953-4419083 | 15 | 8 | 7 | 0 |
|  | 7 | 21.8 Kb | incomplete | 40 | tail, head, integrase | 4924727-4946598 | 16 | 9 | 7 | 0 |
| *Erwinia endophytica* A41C3 | 1 | 8.6 Kb | incomplete | 50 | tail, transposase | 30574-39212 | 10 | 7 | 3 | 0 |
|  | 2 | 49.8 Kb | intact | 140 | tail, head, terminase, lysis, capsid, integrase | 911321-961196 | 72 | 53 | 19 | 9 |
|  | 3 | 8.4 Kb | questionable | 70 | transposase | 989050-997471 | 13 | 8 | 5 | 0 |
|  | 4 | 24.8 Kb | incomplete | 30 | integrase, capsid | 1777219-1802071 | 28 | 18 | 10 | 0 |
|  | 5 | 18.6 Kb | incomplete | 60 | tail, terminase, capsid, integrase | 1796987-1815682 | 18 | 11 | 7 | 0 |
|  | 6 | 18.7 Kb | incomplete | 30 | integrase, tail | 2562480-2581210 | 19 | 13 | 6 | 0 |
|  | 7 | 28.6 Kb | intact | 150 | integrase, transposase, capsid, head, plate, tail | 3038367-3067041 | 28 | 25 | 3 | o |
|  | 8 | 8.4 Kb | incomplete | 30 | flippase, transposase | 3200409-3208835 | 10 | 6 | 4 | 0 |
|  | 9 | 6.3 Kb | questionable | 70 | portal, transposase | 3636109-3642494 | 18 | 11 | 7 | 0 |
|  | 10 | 16.7 Kb | incomplete | 30 | integrase | 4029570-4046350 | 6 | 6 | 0 | 0 |
| *Erwinia mallotivora* ICMP5705^T^ | 1 | 11.9 Kb | incomplete | 30 | integrase, tail | 367523-379484 | 6 | 6 | 0 | 0 |
|  | 2 | 8.2 Kb | incomplete | 30 | transposase | 697132-705337 | 13 | 7 | 6 | 0 |
|  | 3 | 19.2 Kb | incomplete | 30 | transposase | 941747-961041 | 11 | 7 | 4 | 0 |
|  | 4 | 9.5 Kb | incomplete | 40 | integrase, tail | 1573943-1583507 | 14 | 10 | 4 | 0 |
|  | 5 | 21.2 Kb | intact | 140 | protease, tail, plate, capsid, portal, transposase | 1963052-1984331 | 26 | 19 | 7 | 0 |
|  | 6 | 69.3 Kb | questionable | 90 | protease, head, tail, plate | 2210350-2279653 | 77 | 46 | 31 | 0 |
| *Erwinia oleae* DAPP-PG531^T^ | 1 | 10.1 Kb | incomplete | 30 | virion, head | 820062-830247 | 16 | 7 | 9 | 0 |
|  | 2 | 17.3 Kb | incomplete | 30 | tail, integrase | 2488064-2505460 | 18 | 13 | 5 | 0 |
|  | 3 | 14.7 Kb | intact | 100 | transposase, lysin, plate, tail | 2641037-2655807 | 16 | 12 | 4 | 0 |
|  | 4 | 6.2 Kb | incomplete | 40 | transposase | 4137285-4143583 | 7 | 6 | 1 | 0 |
|  | 5 | 14.4 Kb | incomplete | 40 | tail, transposase | 4496670-4511099 | 17 | 10 | 7 | 0 |
| *Erwinia papayae*  JGD 233 | 1 | 68.4 Kb | intact | 150 | tail, head, portal, terminase, integrase | 3099911-3168323 | 61 | 44 | 17 | 0 |
|  | 2 | 11.5 Kb | incomplete | 20 | tail | 4810154-4821677 | 23 | 10 | 13 | 0 |
|  | 3 | 8.3 Kb | incomplete | 30 | tail, transposase | 4882441-4890773 | 10 | 10 | 0 | 0 |
| *Erwinia persicina*  CFBP8797 | 1 | 16.8 Kb | intact | 130 | tail, plate, lysis, lysin | 2805505-2822304 | 22 | 20 | 2 | 0 |
|  | 2 | 16.2 Kb | incomplete | 40 | transposase, integrase, tail | 3310706-3326980 | 13 | 12 | 1 | 0 |
| *Erwinia persicina*  NBRC 102418^T^ | 1 | 18.1 Kb | incomplete | 50 | transposase, integrase | 2150607-2168736 | 10 | 7 | 3 | 0 |
|  | 2 | 16.8 Kb | intact | 140 | tail, plate, lysis, lysin | 2759674-2776544 | 24 | 21 | 3 | 0 |
|  | 3 | 12.2 Kb | incomplete | 50 | transposase, integrase, tail, terminase | 4312885-4325089 | 10 | 6 | 4 | 0 |
|  | 4 | 62.1 Kb | intact | 150 | integrase, transposase, tail, terminase, head, portal | 4485178-4547347 | 76 | 55 | 21 | 0 |
| *Erwinia phyllosphaerae*  CMYE1^T^ | 1 | 49.4 Kb | questionable | 90 | integrase, lysin, tail, terminase | 1023156-1072584 | 68 | 43 | 25 | 0 |
|  | 2 | 56.2 Kb | incomplete | 40 | head | 1397596-1453824 | 67 | 42 | 25 | 0 |
|  | 3 | 15.8 Kb | incomplete | 30 | integrase, tail | 1492362-1508208 | 17 | 11 | 6 | 0 |
|  | 4 | 17.1 Kb | incomplete | 30 | integrase, tail | 3248852-3265953 | 14 | 12 | 2 | 0 |
|  | 5 | 18.7 Kb | incomplete | 60 | lysis, tail, plate | 3272105-3290829 | 22 | 21 | 1 | 0 |
|  | 6 | 40.8 Kb | questionable | 87 | lysin | 3535484-3576309 | 55 | 43 | 12 | 0 |
| *Erwinia piriflorinigrans*  CFBP5888^T^ | 1 | 10.1 Kb | incomplete | 30 | transposase | 439244-449380 | 10 | 6 | 4 | 0 |
|  | 2 | 13 Kb | incomplete | 30 | tail, transposase | 2821072-2834163 | 16 | 12 | 4 | 0 |
|  | 3 | 36.4 Kb | intact | 150 | tail, plate, transposase | 3222245-3258687 | 54 | 34 | 20 | 0 |
| *Erwinia plantamica*  OPT-41^T^ | 1 | 36.5 Kb | intact | 150 | capsid, tail, plate, virion, lysis, head, terminase, portal, integrase | 1481582-1518103 | 46 | 40 | 6 | 0 |
|  | 2 | 16.8 Kb | incomplete | 50 | integrase, tail | 1791047-1807933 | 24 | 9 | 15 | 0 |
|  | 3 | 21.6 Kb | incomplete | 20 | integrase | 2370471-2392113 | 14 | 7 | 7 | 0 |
|  | 4 | 14.5 Kb | incomplete | 20 | integrase, tail | 2452245-2466839 | 19 | 12 | 7 | 0 |
|  | 5 | 20.4 Kb | incomplete | 60 | lysin, portal, capsid, injection, tail | 2469637-2490078 | 26 | 21 | 5 | 0 |
|  | 6 | 23.2 kB | questionable | 90 | integrase, head, transposase, lysis, tail, plate | 3258059-3281272 | 22 | 19 | 3 | 0 |
|  | 7 | 15.4 Kb | incomplete | 20 | tail | 3984292-3999782 | 20 | 14 | 6 | 0 |
| *Erwinia psidii*  IBSBF435^T^ | 1 | 31.1 Kb | intact | 92 | NA | 1115752-1146916 | 34 | 31 | 3 | 0 |
|  | 2 | 7.8 Kb | incomplete | 30 | transposase, tail | 2207277-2215103 | 8 | 7 | 1 | 0 |
|  | 3 | 7.7 Kb | incomplete | 50 | plate, transposase | 2476973-2484735 | 11 | 6 | 5 | 0 |
|  | 4 | 18.7 Kb | incomplete | 30 | integrase, tail | 3027587-3046348 | 19 | 9 | 10 | 0 |
|  | 5 | 36.2 Kb | intact | 96 | lysin | 3501407-3537701 | 50 | 40 | 10 | 0 |
|  | 6 | 18.7 Kb | incomplete | 30 | integrase, tail | 3660280-3679020 | 12 | 11 | 1 | 0 |
|  | 7 | 59.4 Kb | intact | 150 | integrase, portal, terminase, capsid, head, lysis, tail, virion, plate, recombinase | 3789756-3849156 | 67 | 45 | 22 | 0 |
| *Erwinia pyri*  DE2^T^ | 1 | 15.6 Kb | incomplete | 40 | integrase, head | 604268-619886 | 13 | 8 | 5 | 0 |
|  | 2 | 42.9 Kb | incomplete | 30 | tail | 871895-914876 | 19 | 18 | 1 | 0 |
|  | 3 | 16.7 Kb | incomplete | 20 | tail | 1144648-1161352 | 20 | 14 | 6 | 0 |
|  | 4 | 11.4 Kb | incomplete | 30 | head, tail | 1811488-1822919 | 18 | 9 | 9 | 0 |
|  | 5 | 10.2 Kb | incomplete | 30 | lysis | 2518956-2529186 | 16 | 13 | 3 | 0 |
|  | 6 | 42 Kb | intact | 150 | integrase, tail, plate, head, portal, terminase, lysin | 2565799-2607839 | 59 | 50 | 9 | 0 |
|  | 7 | 72.5 Kb | intact | 120 | transposase, portal, terminase, capsid, tail, integrase | 2708391-2780924 | 73 | 57 | 16 | 0 |
|  | 8 | 21.5 Kb | incomplete | 10 | NA | 4377300-4398870 | 30 | 20 | 10 | 0 |
| *Erwinia pyrifoliae*  DSM12163^T^ | 1 | 8.2 Kb | incomplete | 40 | tail, transposase | 685560-693786 | 12 | 8 | 4 | 0 |
|  | 2 | 9.8 Kb | incomplete | 20 | tail | 1059308-1069187 | 13 | 9 | 4 | 0 |
|  | 3 | 12.8 Kb | incomplete | 30 | transposase, tail | 1130588-1143468 | 14 | 12 | 2 | 0 |
|  | 4 | 28.2 Kb | incomplete | 50 | tail, portal | 1977987-2006212 | 44 | 37 | 7 | 0 |
|  | 5 | 45.2 Kb | intact | 150 | head, tail, plate, portal, terminase, integrase | head, tail, plate, portal, terminase, integrase | 63 | 49 | 14 | 0 |
|  | 6 | 23.1 Kb | incomplete | 50 | integrase, transposase | 3201176-3224293 | 21 | 8 | 13 | 0 |
|  | 7 | 11.6 Kb | intact | 150 | transposase | 3494261-3505947 | 23 | 16 | 7 | 0 |
| *Erwinia rhapontici*  BY21311 | 1 | 35.5 Kb | incomplete | 40 | tail, integrase | 3783114-3818633 | 21 | 15 | 6 | 0 |
|  | 2 | 16.4 Kb | intact | 120 | tail, plate, lysis | 4385528-4401979 | 23 | 21 | 2 | 0 |
|  | 3 | 47.1 Kb | intact | 150 | tail, plate, lysin, head, terminase, capsid, portal, integrase | 4880931-4928084 | 50 | 40 | 10 | 0 |
| *Erwinia sorbitola*  J780^T^ | 1 | 27.2 Kb | incomplete | 30 | integrase, tail | 1088921-1116175 | 15 | 12 | 3 | 0 |
|  | 2 | 53.9 Kb | incomplete | 60 | integrase, tail, capsid | 1651990-1705951 | 76 | 60 | 16 | 0 |
| *Erwinia* sp. PL328 | 1 | 9.7 Kb | incomplete | 60 | transposase, tail, integrase | 2048624-2058329 | 12 | 7 | 5 | 0 |
|  | 2 | 13.8 Kb | incomplete | 30 | tail, transposase | 3583319-3597152 | 17 | 14 | 3 | 0 |
|  | 3 | 16.3 Kb | intact | 100 | tail, plate, lysis | 4182516-4198822 | 23 | 21 | 2 | 0 |
|  | 4 | 25.1 Kb | incomplete | 30 | integrase, tail | 4319118-4344301 | 11 | 6 | 5 | 0 |
| *Erwinia tasmaniensis*  Et1/99^T^ | 1 | 35.7 Kb | incomplete | 40 | lysin | 625327-661033 | 44 | 39 | 5 | 0 |
|  | 2 | 28.5 Kb | incomplete | 30 | integrase, tail | 1076115-1104693 | 15 | 12 | 3 | 0 |
| *Erwinia tasmaniensis*  PPS120 | 1 | 14.2 Kb | incomplete | 40 | tail, head | 500240-514502 | 22 | 13 | 9 |  |
|  | 2 | 32.7 Kb | incomplete | 50 | head, integrase, tail | 514102-546893 | 27 | 19 | 8 | 0 |
|  | 3 | 17.2 Kb | incomplete | 60 | tail, plate, transposase | 556157-573439 | 25 | 20 | 5 | 0 |
|  | 4 | 10.2 Kb | incomplete | 40 | transposase | 860469-870767 | 17 | 11 | 6 | 0 |
|  | 5 | 69.7 Kb | intact | 140 | tail, lysis, portal, head, capsid, integrase | 1610845-1680572 | 59 | 36 | 23 | 0 |
|  | 6 | 45.1 Kb | intact | 150 | integrase, portal, terminase, capsid, head, lysis, tail, virion, plate | 3171911-3217094 | 49 | 40 | 9 | 0 |
|  | 7 | 47.6 Kb | intact | 96 | plate, protease | 3444032-3491646 | 50 | 36 | 14 | 0 |
|  | 8 | 11.4 Kb | questionable | 70 | head, transposase, terminase, integrase | 3652518-3663945 | 18 | 12 | 6 | 0 |
|  | 9 | 3.2 Kb | incomplete | 50 | transposase | 3698956-3702242 | 6 | 6 | 0 | 0 |
|  | 10 | 22.7 Kb | incomplete | 60 | tail, transposase, integrase | 4509292-4531992 | 11 | 8 | 3 | 0 |
|  | 11 | 8.8 Kb | incomplete | 50 | transposase, tail | 4545432-4554278 | 16 | 8 | 8 | 0 |
| *Erwinia tracheiphila*  ICMP5845^T^ | 1 | 29.4Kb | incomplete | 20 | transposase | 91809-121228 | 21 | 7 | 14 | 0 |
|  | 2 | 25.8Kb | incomplete | 40 | plate, transposase, integrase | 175823-201707 | 16 | 10 | 6 | 0 |
|  | 3 | 18.7Kb | incomplete | 30 | transposase | 270023-288788 | 21 | 11 | 10 | 0 |
|  | 4 | 28.4Kb | incomplete | 50 | integrase, transposase | 446533-474999 | 13 | 8 | 5 | 0 |
|  | 5 | 4.7Kb | incomplete | 10 | NA | 751560-756300 | 7 | 6 | 1 | 0 |
|  | 6 | 33.4Kb | intact | 140 | tail, transposase, plate | 765307-798736 | 30 | 21 | 9 | 0 |
|  | 7 | 15.3Kb | intact | 120 | transposase | 872844-888171 | 31 | 17 | 14 | 0 |
|  | 8 | 24.4Kb | incomplete | 50 | protease, transposase | 887326-911804 | 21 | 9 | 12 | 0 |
|  | 9 | 26.1Kb | intact | 150 | transposase, head, integrase | 920236-946359 | 29 | 18 | 11 | 0 |
|  | 10 | 49.1Kb | intact | 150 | transposase, portal, virion, protease, head, tail, plate, recombinase | 1005076- 1054205 | 66 | 48 | 18 | 0 |
|  | 11 | 6.1Kb | incomplete | 20 | plate | 1220695-1226839 | 9 | 6 | 3 | 0 |
|  | 12 | 42.3Kb | intact | 100 | lysin | 1361492-1403857 | 75 | 52 | 23 | 0 |
|  | 13 | 13.1Kb | incomplete | 30 | integrase, transposase | 1407932-1421055 | 12 | 6 | 6 | 0 |
|  | 14 | 13.3Kb | intact | 110 | transposase | 1542906-1556226 | 25 | 15 | 10 | 0 |
|  | 15 | 25.5Kb | questionable | 70 | transposase, integrase, tail | 1747717-1773274 | 24 | 14 | 10 | 0 |
|  | 16 | 29.2Kb | intact | 120 | lysin | 1768923-1798200 | 45 | 45 | 0 | 0 |
|  | 17 | 38Kb | intact | 150 | transposase, integrase, tail, terminase | 1848448-1886469 | 51 | 29 | 22 | 0 |
|  | 18 | 18Kb | incomplete | 50 | transposase, integrase | 1973899-1991925 | 11 | 8 | 3 | 0 |
|  | 19 | 31Kb | intact | 150 | transposase, tail, plate, head, protease, virion, portal | 2128127-2159212 | 65 | 44 | 21 | 0 |
|  | 20 | 18.2Kb | questionable | 90 | transposase, integrase | 2350290-2368526 | 31 | 19 | 12 | 0 |
|  | 21 | 62Kb | intact | 150 | transposase, integrase, capsid, tail, head, portal, lysis, terminase, injection | 2370656-2432717 | 100 | 72 | 28 | 0 |
|  | 22 | 36.4Kb | intact | 140 | tail, transposase, lysin | 2460781-2497233 | 54 | 48 | 6 | 0 |
|  | 23 | 8Kb | intact | 100 | virion, transposase | 3051536-3059599 | 15 | 10 | 5 | 0 |
|  | 24 | 35.1Kb | intact | 150 | terminase, transposase | 3059640-3094761 | 35 | 20 | 15 | 0 |
|  | 25 | 10.3Kb | intact | 150 | capsid, transposase | 3173958-3184298 | 26 | 14 | 12 | 0 |
|  | 26 | 27.3Kb | incomplete | 40 | integrase, transposase | 3188947-3216334 | 15 | 8 | 7 | 0 |
|  | 27 | 9.2Kb | questionable | 70 | tail, transposase | 3218141-3227379 | 22 | 11 | 11 | 0 |
|  | 28 | 33.5Kb | intact | 120 | head, transposase | 3250876-3284418 | 21 | 12 | 9 | 0 |
|  | 29 | 22Kb | intact | 150 | capsid, integrase | 3281484-3303539 | 44 | 23 | 21 | 0 |
|  | 30 | 19.9Kb | intact | 150 | head, transposase, tail | 3319658-3339611 | 38 | 20 | 18 | 0 |
|  | 31 | 11.8Kb | intact | 100 | integrase, transposase | 3403923-3415736 | 22 | 12 | 10 | 0 |
|  | 32 | 15.6Kb | incomplete | 40 | tail, protease | 3652792-3668488 | 28 | 15 | 13 | 0 |
|  | 33 | 14.7Kb | questionable | 90 | transposase | 3742844-3757557 | 32 | 17 | 15 | 0 |
|  | 34 | 70.1Kb | intact | 150 | capsid, head, tail, integrase | 3752208-3822359 | 101 | 60 | 41 | 0 |
|  | 35 | 10.3Kb | intact | 100 | portal, transposase | 3838007-3848317 | 22 | 11 | 11 | 0 |
|  | 36 | 52.1Kb | intact | 150 | terminase, head, tail | 3887764-3939955 | 68 | 39 | 29 | 0 |
|  | 37 | 40.4Kb | questionable | 88 | protease, integrase, transposase | 3984149-4024584 | 56 | 28 | 28 | 0 |
|  | 38 | 10.1Kb | intact | 100 | capsid, transposase | 4033606-4043776 | 18 | 9 | 9 | 0 |
|  | 39 | 9Kb | incomplete | 60 | integrase, tail | 4070912-4080001 | 13 | 7 | 6 | 0 |
|  | 40 | 31.5Kb | intact | 130 | portal, head, transposase | 4187651-4219236 | 39 | 22 | 17 | 0 |
|  | 41 | 16.3Kb | incomplete | 60 | tail, protease | 4215292-4231638 | 28 | 15 | 13 | 0 |
|  | 42 | 47.1Kb | intact | 150 | terminase, head, tail, integrase | 4251084-4298213 | 76 | 40 | 36 | 0 |
|  | 43 | 15.2Kb | questionable | 70 | transposase | 4309561-4324776 | 28 | 15 | 13 | 0 |
|  | 44 | 29.3Kb | intact | 150 | portal, plate, head, transposase | 4572774-4602096 | 53 | 29 | 24 | 0 |
|  | 45 | 9.5Kb | questionable | 70 | integrase | 4680434-4690023 | 21 | 11 | 10 | 0 |
|  | 46 | 30.7Kb | incomplete | 30 | transposase | 4692494-4723209 | 15 | 7 | 8 | 0 |
|  | 47 | 38.4Kb | incomplete | 50 | tail, head, transposase | 4848347-4886789 | 30 | 16 | 14 | 0 |
| *Erwinia typographi*  M043b | 1 | 15.9 Kb | incomplete | 50 | capsid, transposase, tail, integrase | 545509-561443 | 10 | 7 | 3 | 0 |
|  | 2 | 33.1 Kb | incomplete | 50 | tail, integrase | 923065-956164 | 39 | 36 | 3 | 0 |
|  | 3 | 36.2 Kb | intact | 150 | tail, plate, head, virion, portal, terminase, transposase | 963295-999561 | 56 | 37 | 19 | 0 |
|  | 4 | 8.5 Kb | incomplete | 50 | transposase, tail, plate, protease | 1283566-1292113 | 15 | 8 | 7 | 0 |
|  | 5 | 19.4 Kb | incomplete | 30 | integrase, transposase | 2208097-2227567 | 10 | 6 | 4 | 0 |
|  | 6 | 27.4 Kb | incomplete | 30 | tail, integrase | 3007590-3035010 | 14 | 12 | 2 | 0 |
|  | 7 | 12.4 Kb | incomplete | 10 | NA | 3603701-3616164 | 22 | 11 | 11 | 0 |
|  | 8 | 12.2 Kb | incomplete | 20 | lysin | 4189052-4201348 | 16 | 9 | 7 | 0 |
|  | 9 | 16.3 Kb | questionable | 90 | terminase, head, portal, capsid, integrase | 4514924-4531290 | 21 | 13 | 8 | 0 |
|  | 10 | 8.6 Kb | questionable | 80 | capsid, tail | 4746760-4755381 | 11 | 7 | 4 | 0 |
|  | 11 | 13.9 Kb | incomplete | 40 | flippase, tail, integrase | 4752348-4766257 | 11 | 6 | 5 | 0 |
|  | 12 | 44 Kb | intact | 150 | head, lysin, terminase, portal, protease, tail | 5013123-5057124 | 51 | 43 | 8 | 0 |
|  | 13 | 7 Kb | incomplete | 30 | transposase, capsid | 5086587-5093640 | 9 | 6 | 3 | 0 |
|  | 14 | 9.3 Kb | incomplete | 50 | transposase, tail | 5183765-5193077 | 14 | 8 | 6 | 0 |
|  | 15 | 23.2 Kb | intact | 100 | tail, portal, head, terminase | 5536758-5560054 | 31 | 23 | 8 | 0 |
|  | 16 | 41.4 Kb | intact | 150 | transposase, integrase | 5663297-5704747 | 27 | 15 | 12 | 0 |
|  | 17 | 8.1 Kb | incomplete | 60 | transposase, tail | 5720036-5728164 | 10 | 8 | 2 | 0 |
