## Supplementary for "Phylogenomics and comparative genomics of the genus *Erwinia* reveal taxonomic inconsistencies and evolutionary diversification": Supplementary Table 5.docx

**Supplementary Table 5.** Antimicrobial Resistance genes identified in *Erwinia persicina* CFBP8797 and *Erwinia persicina* NBRC 102418 ^T^

| **Antimicrobial** | **Class** | ***Erwinia persicina* CFBP8797** | | ***Erwinia persicina* NBRC 102418^T^** | |
| --- | --- | --- | --- | --- | --- |
|  |  | **WGS-predicted phenotype** | **Match^*^** | **WGS-predicted phenotype** | **Match^*^** |
| gentamicin | aminoglycoside | No resistance | 0 | No resistance | 0 |
| tobramycin | aminoglycoside | No resistance | 0 | No resistance | 0 |
| streptomycin | aminoglycoside | No resistance | 0 | No resistance | 0 |
| amikacin | aminoglycoside | No resistance | 0 | No resistance | 0 |
| isepamicin | aminoglycoside | No resistance | 0 | No resistance | 0 |
| dibekacin | aminoglycoside | No resistance | 0 | No resistance | 0 |
| kanamycin | aminoglycoside | No resistance | 0 | No resistance | 0 |
| neomycin | aminoglycoside | No resistance | 0 | No resistance | 0 |
| lividomycin | aminoglycoside | No resistance | 0 | No resistance | 0 |
| paromomycin | aminoglycoside | No resistance | 0 | No resistance | 0 |
| ribostamycin | aminoglycoside | No resistance | 0 | No resistance | 0 |
| unknown aminoglycoside | aminoglycoside | No resistance | 0 | No resistance | 0 |
| butiromycin | aminoglycoside | No resistance | 0 | No resistance | 0 |
| butirosin | aminoglycoside | No resistance | 0 | No resistance | 0 |
| hygromycin | aminoglycoside | No resistance | 0 | No resistance | 0 |
| netilmicin | aminoglycoside | No resistance | 0 | No resistance | 0 |
| apramycin | aminoglycoside | No resistance | 0 | No resistance | 0 |
| sisomicin | aminoglycoside | No resistance | 0 | No resistance | 0 |
| arbekacin | aminoglycoside | No resistance | 0 | No resistance | 0 |
| kasugamycin | aminoglycoside | No resistance | 0 | No resistance | 0 |
| astromicin | aminoglycoside | No resistance | 0 | No resistance | 0 |
| fortimicin | aminoglycoside | No resistance | 0 | No resistance | 0 |
| spectinomycin | aminocyclitol | No resistance | 0 | No resistance | 0 |
| fluoroquinolone | quinolone | No resistance | 0 | No resistance | 0 |
| ciprofloxacin | quinolone | No resistance | 0 | No resistance | 0 |
| unknown quinolone | quinolone | No resistance | 0 | No resistance | 0 |
| nalidixic acid | quinolone | No resistance | 0 | No resistance | 0 |
| **amoxicillin** | **beta-lactam** | **Resistant** | **2** | **Resistant** | **3** |
| amoxicillin+clavulanic acid | beta-lactam | No resistance | 0 | No resistance | 0 |
| **ampicillin** | **beta-lactam** | **Resistant** | **2** | **Resistant** | **3** |
| ampicillin+clavulanic acid | beta-lactam | No resistance | 0 | No resistance | 0 |
| cefepime | beta-lactam | No resistance | 0 | No resistance | 0 |
| cefixime | beta-lactam | No resistance | 0 | No resistance | 0 |
| cefotaxime | beta-lactam | No resistance | 0 | No resistance | 0 |
| cefoxitin | beta-lactam | No resistance | 0 | No resistance | 0 |
| ceftazidime | beta-lactam | No resistance | 0 | No resistance | 0 |
| ertapenem | beta-lactam | No resistance | 0 | No resistance | 0 |
| imipenem | beta-lactam | No resistance | 0 | No resistance | 0 |
| meropenem | beta-lactam | No resistance | 0 | No resistance | 0 |
| **piperacillin** | **beta-lactam** | **Resistant** | **2** | **Resistant** | **3** |
| piperacillin+tazobactam | beta-lactam | No resistance | 0 | No resistance | 0 |
| unknown beta-lactam | beta-lactam | No resistance | 0 | No resistance | 0 |
| aztreonam | beta-lactam | No resistance | 0 | No resistance | 0 |
| cefotaxime+clavulanic acid | beta-lactam | No resistance | 0 | No resistance | 0 |
| temocillin | beta-lactam | No resistance | 0 | No resistance | 0 |
| **ticarcillin** | **beta-lactam** | **Resistant** | **2** | **Resistant** | **3** |
| ceftazidime+avibactam | beta-lactam | No resistance | 0 | No resistance | 0 |
| penicillin | beta-lactam | No resistance | 0 | No resistance | 0 |
| ceftriaxone | beta-lactam | No resistance | 0 | No resistance | 0 |
| ticarcillin+clavulanic acid | beta-lactam | No resistance | 0 | No resistance | 0 |
| cephalothin | beta-lactam | No resistance | 0 | No resistance | 0 |
| piperacillin+clavulanic acid | beta-lactam | No resistance | 0 | No resistance | 0 |
| ceftiofur | under_development | No resistance | 0 | No resistance | 0 |
| sulfamethoxazole | folate pathway antagonist | No resistance | 0 | No resistance | 0 |
| trimethoprim | folate pathway antagonist | No resistance | 0 | No resistance | 0 |
| fosfomycin | fosfomycin | No resistance | 0 | No resistance | 0 |
| vancomycin | glycopeptide | No resistance | 0 | No resistance | 0 |
| teicoplanin | glycopeptide | No resistance | 0 | No resistance | 0 |
| bleomycin | glycopeptide | No resistance | 0 | No resistance | 0 |
| lincomycin | lincosamide | No resistance | 0 | No resistance | 0 |
| clindamycin | lincosamide | No resistance | 0 | No resistance | 0 |
| dalfopristin | streptogramin a | No resistance | 0 | No resistance | 0 |
| pristinamycin iia | streptogramin a | No resistance | 0 | No resistance | 0 |
| virginiamycin m | streptogramin a | No resistance | 0 | No resistance | 0 |
| quinupristin+dalfopristin | streptogramin a | No resistance | 0 | No resistance | 0 |
| tiamulin | pleuromutilin | No resistance | 0 | No resistance | 0 |
| carbomycin | macrolide | No resistance | 0 | No resistance | 0 |
| erythromycin | macrolide | No resistance | 0 | No resistance | 0 |
| azithromycin | macrolide | No resistance | 0 | No resistance | 0 |
| oleandomycin | macrolide | No resistance | 0 | No resistance | 0 |
| spiramycin | macrolide | No resistance | 0 | No resistance | 0 |
| tylosin | macrolide | No resistance | 0 | No resistance | 0 |
| telithromycin | macrolide | No resistance | 0 | No resistance | 0 |
| tetracycline | tetracycline | No resistance | 0 | No resistance | 0 |
| doxycycline | tetracycline | No resistance | 0 | No resistance | 0 |
| minocycline | tetracycline | No resistance | 0 | No resistance | 0 |
| tigecycline | tetracycline | No resistance | 0 | No resistance | 0 |
| quinupristin | streptogramin b | No resistance | 0 | No resistance | 0 |
| pristinamycin ia | streptogramin b | No resistance | 0 | No resistance | 0 |
| virginiamycin s | streptogramin b | No resistance | 0 | No resistance | 0 |
| linezolid | oxazolidinone | No resistance | 0 | No resistance | 0 |
| chloramphenicol | amphenicol | No resistance | 0 | No resistance | 0 |
| florfenicol | amphenicol | No resistance | 0 | No resistance | 0 |
| colistin | polymyxin | No resistance | 0 | No resistance | 0 |
| fusidic acid | steroid antibacterial | No resistance | 0 | No resistance | 0 |
| mupirocin | pseudomonic acid | No resistance | 0 | No resistance | 0 |
| rifampicin | rifamycin | No resistance | 0 | No resistance | 0 |
| metronidazole | nitroimidazole | No resistance | 0 | No resistance | 0 |
| narasin | ionophores | No resistance | 0 | No resistance | 0 |
| salinomycin | ionophores | No resistance | 0 | No resistance | 0 |
| maduramicin | ionophores | No resistance | 0 | No resistance | 0 |

***** The 'Match' column stores one of the integers 0, 1, 2, 3:

**0:** No match found, **1:** Match < 100% ID and match length < ref length, Match = 100% ID and match length < ref length, **3:** Match = 100% ID and match length = ref length
