## Supplementary figures and images for "Phylogenomics and comparative genomics of the genus *Erwinia* reveal taxonomic inconsistencies and evolutionary diversification"

### Supplementary Figure 1.tif

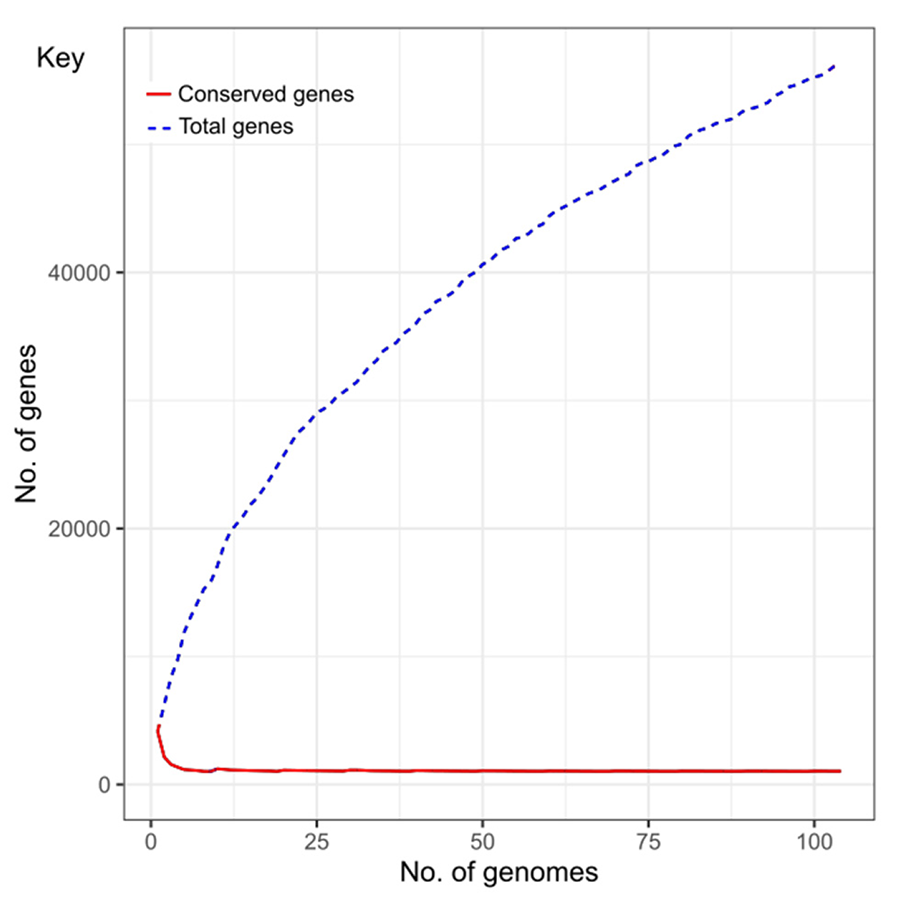

### Supplementary Figure 2.tif

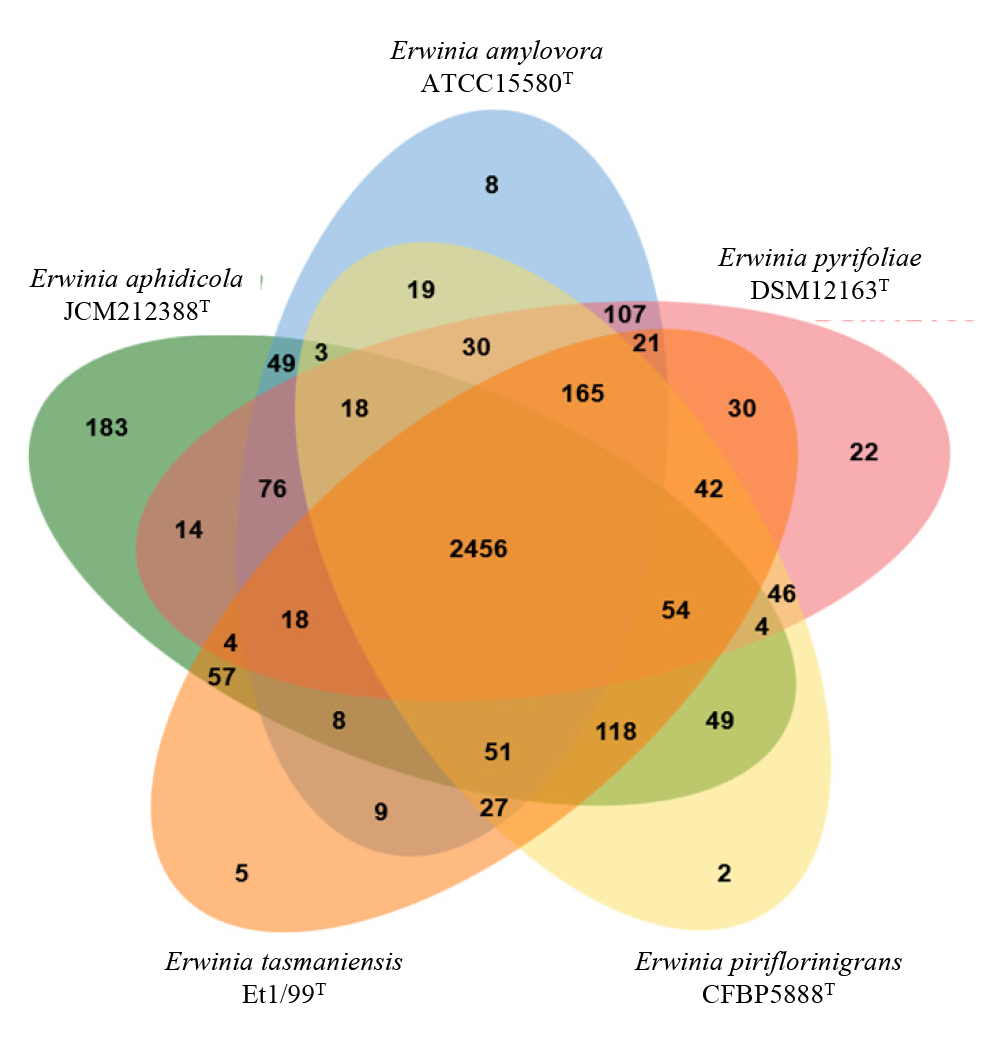

### Supplementary Figure 3.tif

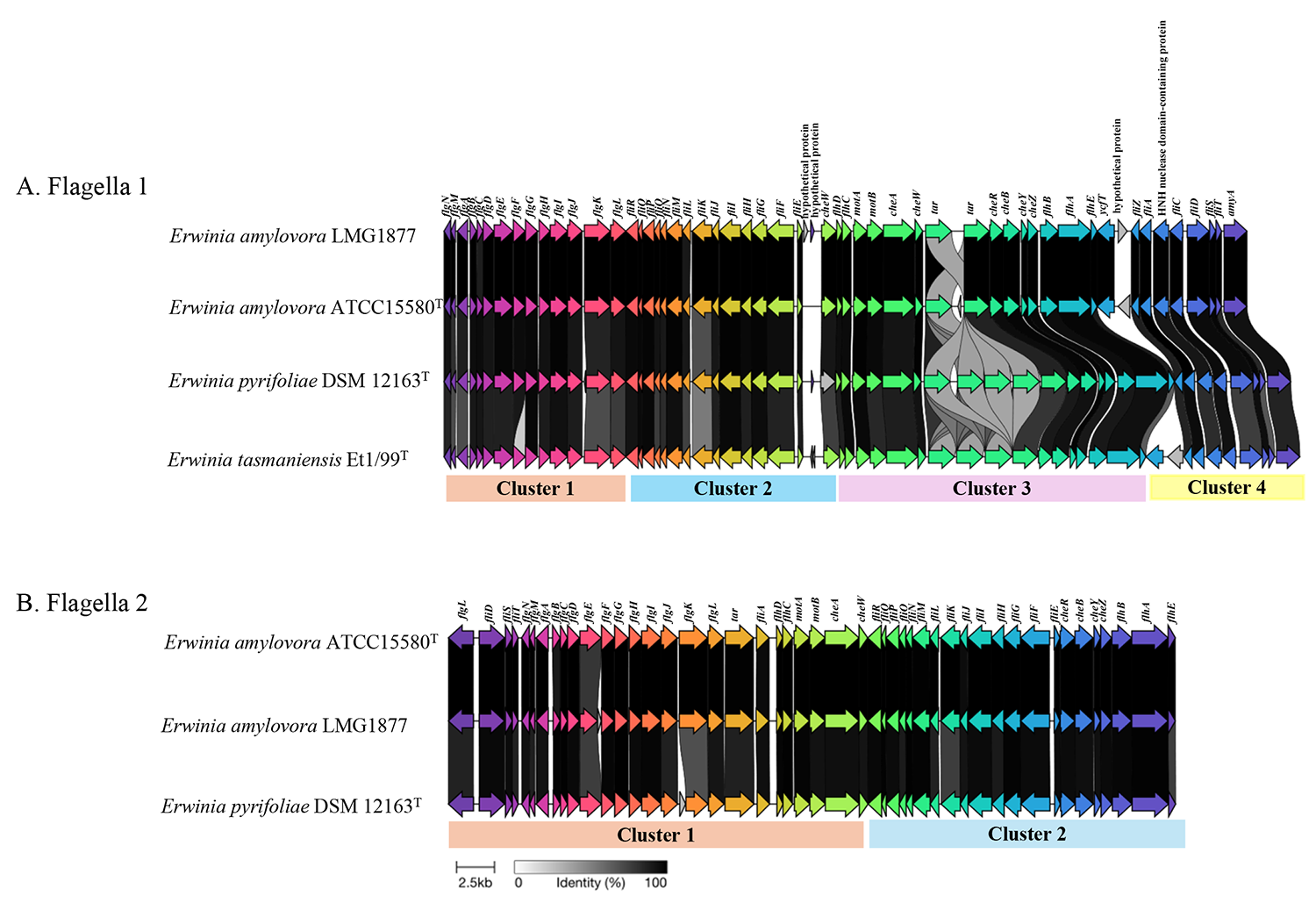
